# Cell-to-cell transmission of HSV1 in human keratinocytes in the absence of the major entry receptor, nectin1

**DOI:** 10.1101/2021.05.13.443971

**Authors:** Joanne Kite, Tiffany Russell, Juliet Jones, Gillian Elliott

## Abstract

Herpes simplex virus 1 (HSV1) infects the stratified epithelia of the epidermis, oral or genital mucosa, where the main cell type is the keratinocyte. Here we have used nTERT human keratinocytes to generate a CRISPR-Cas9 knockout (KO) of the primary candidate HSV1 receptor, nectin1, resulting in a cell line that is refractory to HSV1 entry. Nonetheless, a small population of KO cells was able to support infection, and strikingly at later times, those cells originally resistant to HSV1 infection had also been infected. Appearance of this later population was blocked by inhibition of virus genome replication, or infection with a *Δ*UL34 virus defective in capsid export to the cytoplasm, while newly formed GFP-tagged capsids were detected in cells surrounding the initial infected cell, suggesting that virus was spreading following replication in the original susceptible cells. Neutralizing human serum failed to block virus transmission in KO cells, which was dependent on the cell-to-cell spread glycoproteins gI and gE, indicating that virus was spreading by direct cell-to-cell transmission. In line with these results, both HSV1 and HSV2 formed plaques on nectin1 KO cells, albeit at a reduced titre, confirming that once the original cell population was infected, the virus could spread into all other cells in the monolayer. Additional siRNA depletion of the second major HSV1 receptor HVEM, or PTP1B, a cellular factor shown elsewhere to be involved in cell-to-cell transmission, had no effect on virus spread in the absence of nectin1. We conclude that although nectin1 is required for extracellular entry in to the majority of human keratinocytes, it is dispensable for direct cell-to-cell transmission.

**Author Summary:** Herpes simplex virus 1 (HSV1) infects the epithelia of the epidermis, oral or genital mucosa to cause cold sores, genital herpes, or more serious outcomes such as keratitis and neonatal herpes. Like many viruses, HSV1 can spread through the extracellular environment or by direct cell-to-cell transmission, with the latter mechanism being important for avoiding antibody responses in the host. Here we have studied HSV1 entry and transmission in the human keratinocyte, the main cell type in the target epithelia, by generating a CRISPR-Cas9 knockout of the primary candidate virus receptor, nectin1. While HSV1 was unable to infect the majority of nectin1 knockout keratinocytes, a small population of these nectin1 KO cells remained susceptible to virus entry, and once infected, the virus was able to spread unhindered into the rest of the monolayer. This spread continued in the presence of neutralising serum which blocks extracellular virus, and required the glycoproteins gE and gI which are known to be involved in cell-to-cell spread. Hence, while nectin1 is required for virus entry into the majority of human keratinocyte cells, it is dispensable for cell-to-cell transmission of the virus. These data have implications for the mechanism of HSV1 epithelial spread and pathogenesis.

## Introduction

Herpes simplex virus type 1(HSV1) infects and is generally restricted to the stratified epithelial cells of the skin, oral mucosa, cornea or the genital mucosa before transmitting to and establishing lifelong latent infection in sensory neurons [1]. It reactivates periodically to re-enter the infectious cycle, when it travels back to the epithelia at the initial site of infection and replicates to cause either the cold sore, ocular herpes or genital herpes. Given that the major cell type of these epithelia is the keratinocyte, the molecular events involved in HSV entry and transmission in these cells are highly relevant to understanding the biology of the virus. The most frequently used keratinocyte cell line in HSV1 studies to date has been the HaCaT cell line, a spontaneously immortalised aneuploid line that is unable to undergo terminal epidermal differentiation [2]. By contrast, we have been studying HSV1 entry in the nTERT keratinocyte line, a diploid line derived from primary human keratinocytes. These cells have been retrovirally induced to express the catalytic subunit of the telomerase holoenzyme hTERT, and have spontaneously lost the function of p16^INK4a^ which usually inhibits the transition from G1 to S phase, allowing the maintenance of growth for much longer than normal primary cell types [3]. Crucially, nTERT cells are still dependent on epidermal growth factor and are able to differentiate [4]. Hence, these cells offer a tractable system that are physiologically relevant for the investigation of HSV1 infection of human tissue. Of note, HSV1 enters and replicates rapidly in these human keratinocytes [5], indicating that the virus is exquisitely adapted to grow in these cells.

HSV1 entry involves four essential virus-encoded envelope proteins – the receptor binding protein glycoprotein D (gD), and the core fusion machinery comprising glycoproteins gB, gH and gL [6]. Two major cell receptors have been identified for gD - HveC, more commonly known as the adhesion molecule nectin1 [7], and HVEM (HveA), the first HSV-1 receptor identified, and a member of the tumour necrosis factor receptor family [8] which is expressed predominantly on lymphoid cells [9]. Nectin1 is expressed in both keratinocytes and neurons [7] and has been shown to be highly expressed in HaCaT cells [10]. As such, our previous siRNA depletion studies identifying nectin1 as the preferred receptor over HVEM for HSV1 entry into human nTERT keratinocytes as well as HeLa cells [5], was in broad agreement with other work on human and murine keratinocytes [11]. HSV1 entry is generally studied using cell-free virus infection of monolayers, but cell-to-cell spread of HSV1, whereby virus transmits directly from already infected cells [12], is proposed to be the main route of spread in infected human tissue [13]. Nonetheless, the molecular mechanism of direct cell-to-cell transmission of HSV1 is poorly characterised. The four entry proteins gD, gB, gH and gL are sufficient for infection from the extracellular environment, but efficient infection by cell-to-cell transfer additionally requires glycoproteins gE and gI [14], which are non-essential in tissue culture, but essential for *in vivo* infection [15] adding further weight to the importance of cell-to-cell spread in the host. Although one study has reported that nectin1 is required for cell-to-cell spread in a number of cell types, keratinocytes were not included in that study [16]. Moreover, it was recently suggested that inhibition of the ER-bound protein tyrosine phosphatase 1B (PTP1B) specifically blocks cell-to-cell spread of HSV1, suggesting that tyrosine phosphorylation regulates either the cellular or viral trafficking proteins that make up the cell-to-cell spread machinery [17].

In this current study we wished to generate a tractable tool for HSV1 entry studies by using CRISPR-Cas9 to knock out nectin1 expression in nTERT keratinocytes. As anticipated from previous studies, this resulted in a cell line that was broadly resistant to HSV1 infection, and which lost that resistance when stably transfected to express nectin1 tagged with a V5 epitope. Nonetheless, a small population of nectin1 knockout cells retained susceptibility to HSV1 infection. Intriguingly, these cells supported virus replication, with progeny virus spreading unhindered in the nectin1 KO monolayers, allowing for the production of plaques – a property shared with HSV2. This transmission did not require HVEM or PTP1B, and was not blocked by the presence of neutralising human serum suggesting that it was able to occur by direct cell-to-cell spread. Nonetheless, *Δ*gI or *Δ*gE variants of HSV1, which infected the same proportion of KO cells as Wt virus, were equally defective for virus spread on KO cells and nTERT cells, indicating that this complex was still involved in transmission in the absence of nectin1. Hence, we have discovered that the gI-gE-dependent cell-to-cell transmission pathway of HSV1, which is important for virus spread in the host, can function independently of nectin1, the major HSV1 entry receptor in human keratinocytes.

## Results

### Human keratinocyte cells express high levels of the nectin1 receptor

HSV1 infects keratinocyte cells in the human. It is therefore noteworthy that our previous work has shown that nTERT keratinocyte cells express around 60-fold more nectin1 mRNA than HeLa cells [5]. Moreover, the spontaneously transformed immortal keratinocyte HaCaT cell line also expresses high levels of cell surface nectin1 protein [10]. In our previous work, we did not have suitable tools available to measure endogenous nectin1 protein levels, but assuming this mRNA level were to be translated into protein level similar to HaCaT cells, it would suggest that the natural target cell for the virus is generally extremely rich in its major receptor. In turn, this could explain our previous data demonstrating rapid entry into keratinocytes [5]. Having now acquired an antibody which recognises human nectin1 by flow cytometry, and to a lesser extent immunofluorescence, we have extended this data to determine the level of nectin1 protein on the cell surface of relevant cell types. RT-qPCR confirmed the high level of nectin1 mRNA in nTERT cells compared to HeLa cells (Fig. 1A). Flow cytometry on non-permeabilised cells stained for cell-surface nectin1 revealed that nTERT and HaCaT keratinocytes express around 50-fold and 20-fold more nectin1 respectively on their cell surface compared to HeLa cells (Fig 1B & 1C). When the same antibody was used to stain the surface of each of these cell types for analysis by confocal microscopy, nectin1 was readily detectable on nTERT and HaCaT cells but not on HeLa cells (Fig 1D) confirming that high level cell surface expression of the HSV1 nectin1 receptor is a feature of human keratinocytes. As all these cell types support HSV1 entry, and our previous data suggests that at least in HeLa cells this is via nectin1 [5], we surmise that there is sufficient nectin1 receptor on HeLa cells to support virus entry, but that our previous determination of rapid entry into keratinocyte cells is due at least in part to this much higher availability of the nectin1 receptor for HSV1 [5].

**Figure 1.**
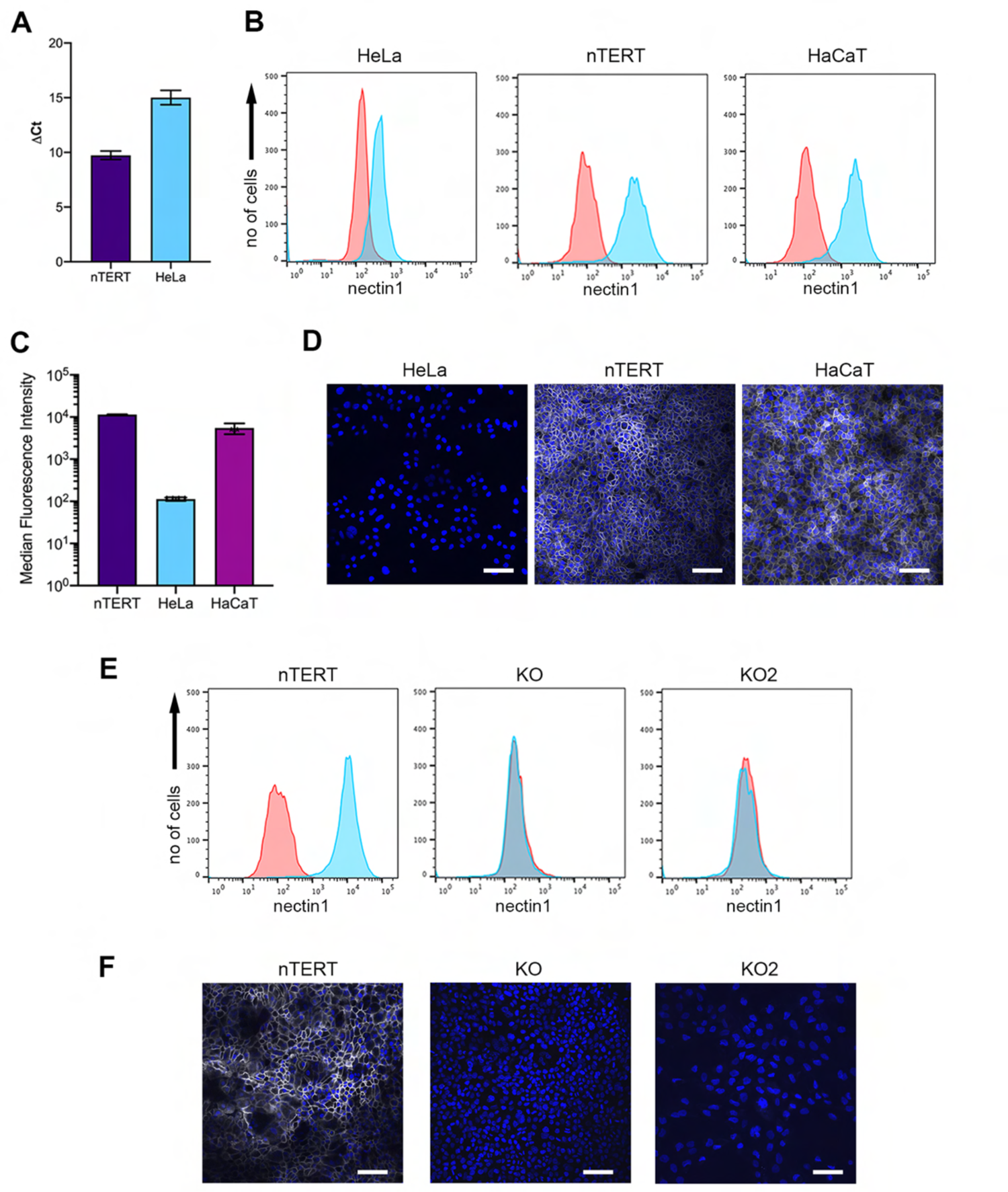
Human keratinocytes express high levels of the major HSV1 receptor, nectin1. **(A)** Total RNA was isolated from HeLa and nTERT cells, and subjected to RT-qPCR using primers for nectin1. Results are presented as *Δ*Ct measurement using 18s as reference (mean±SEM, *n* = 3). **(B)** HeLa, nTERT and HaCaT cells were fixed and stained for nectin1 (blue) or secondary alone (red) before being analysed by flow cytometry. **(C)** Median fluorescence intensity of cell surface nectin1 staining in nTERT, HeLa and HaCaT cells (mean±SEM, *n* = 3). **(D)** HeLa, nTERT and HaCaT cells were grown on coverslips before fixation and staining of non-permeabilised cells for nectin1. Scale bar = 100 μm. **(E)** nTERT, KO and KO2 cells were stained and fixed for nectin1 (blue) or secondary alone (red) before being analysed by flow cytometry. **(F)** nTERT, KO and KO2 cells were grown on coverslips before fixation and staining of non-permeabilised cells for nectin1. Scale bar = 100 μm.

### Generation of a CRISPR-Cas9 nectin1 knockout in nTERT human keratinocytes

To further investigate the contribution of nectin1 to HSV1 entry into keratinocyte cells, and to establish a tractable tool for entry studies, we next generated an nTERT nectin1 knockout cell line using CRISPR-Cas9 mediated gene editing, focusing our efforts on the diploid nTERT cell type rather than the aneuploid HaCaT cell line. nTERT cells transfected with plasmids expressing Cas9 and *NECTIN-1* gRNAs targeting the 5” end of the gene were selected by growth in puromycin, followed by clonal isolation. Candidate nectin1 knockout (KO) cells were cell-surface stained and analysed by flow cytometry and confocal microscopy to determine nectin1 expression on these cells, with only results of the cell line selected for future work being shown here (KO, Fig 1E & 1F). This confirmed that in comparison to nTERT cells these KO cells did not express nectin1 to a level detectable by either flow cytometry or confocal microscopy. Subsequent sequencing of the nectin1 gene revealed that these KO cells were a mixed population containing a number of CRISPR-Cas9 induced edits in the sequences around the gRNA site (not shown) but importantly no parental sequence was present. Due to the nature of their selection, these cells were found to constitutively express Cas9 (not shown), providing a possible explanation for the presence of multiple changes in the nectin1 sequence. Although the phenotype of these cells was clearly nectin1 knockout, we also additionally isolated a partner CRISPR-Cas9 edited cell-line through single cell sorting of transfected cells 72 hours after transfection rather than by puromycin selection, to produce cells lacking both nectin1 and Cas9 expression (KO2, Fig 1E & 1F). In studies presented below, we found no difference in the behaviour of the Cas9 expressing and non-expressing nectin1 knockout cells.

### Characterisation of HSV1 infection in nectin1 KO nTERT cells

For preliminary analysis of the ability of these nectin1 KO cells to support HSV1 entry, cells were synchronously infected with HSV1 expressing *β-galactosidase* under the control of the immediate early (IE) ICP0 promoter [18], and after 3 h, *β-galactosidase* activity was measured as a surrogate for virus entry [19] (Fig 2A). In contrast to nTERT cells, where *β-galactosidase* activity was high, both nectin1 KO lines expressed greatly reduced levels of *β-galactosidase*, suggesting that early events in HSV1 infection had indeed been abrogated.

**Figure 2.**
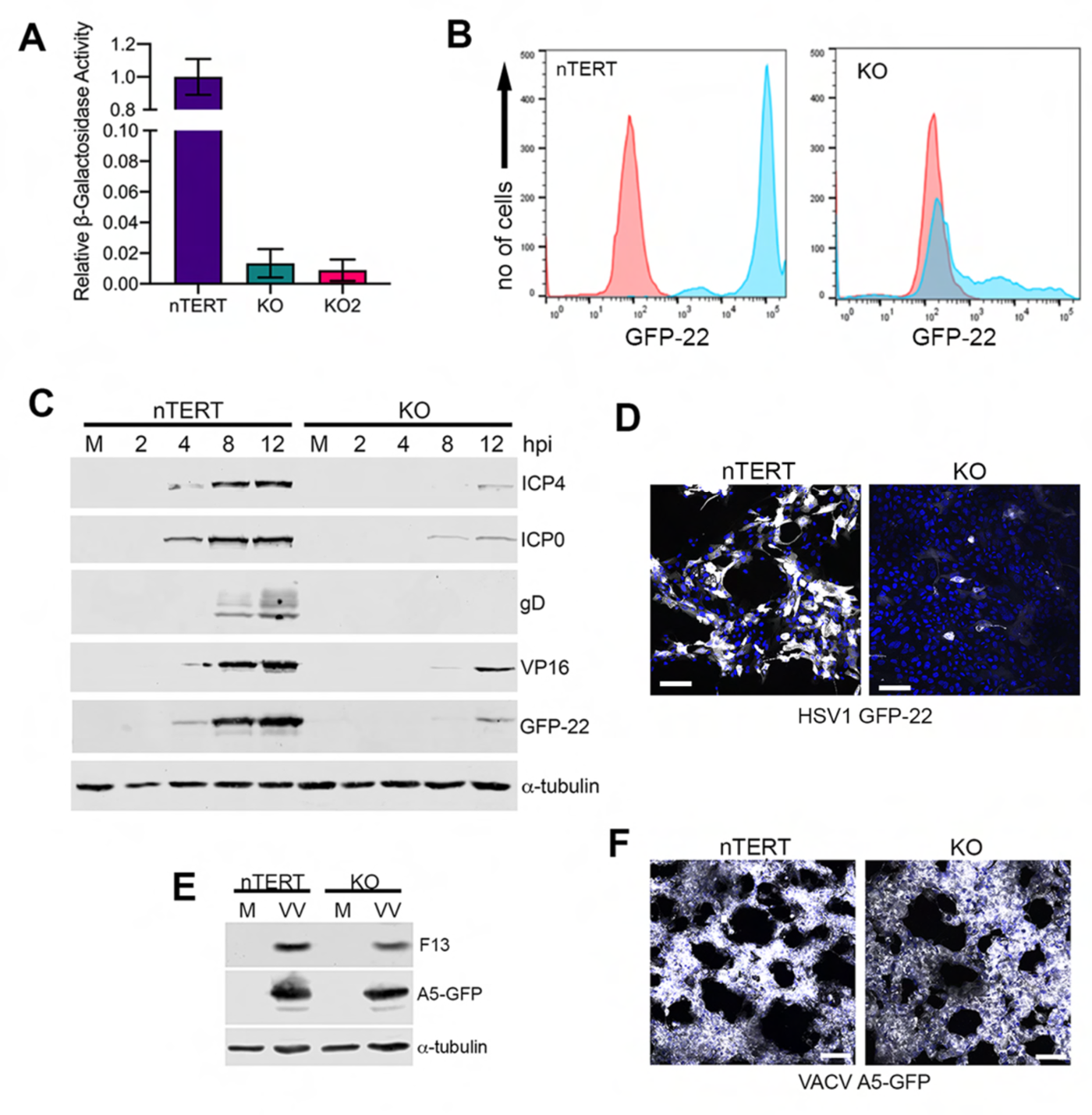
HSV1 infection of nectin1 KO cells. **(A)** nTERT, KO and KO2 cells were infected at MOI 5 with HSV1 IE110*lacZ* on ice to allow attachment, before shifting to 37 °C for 3 h, and β-gal activity was measured by ONPG assay (mean±SEM, *n* = 3). **(B)** nTERT and KO cells were infected with HSV1 expressing GFP-22 at MOI 5, and fixed 8 h later. Uninfected (red) and infected (blue) cells were analysed for GFP fluorescence by flow cytometry. **(C)** nTERT and KO cells were infected with HSV1 GFP-22 at MOI 5, harvested at the indicated times and analysed by SDS-PAGE and Western blotting for a range of virus proteins as indicated, and *α*-tubulin as a loading control. **(D)** nTERT and KO cells grown on coverslips were infected with HSV1 GFP-22 at MOI 5, fixed at 8 h, stained with DAPI (blue) and analysed by confocal microscopy for GFP fluorescence (white). Scale bar = 100 μm. **(E)** nTERT and KO cells were infected with VACV expressing A5GFP at MOI 5, harvested at 16 h and analysed by SDS-PAGE and Western blotting for F13 and GFP. **(F)** nTERT and KO cells grown on coverslips were infected with VACV as in (E), fixed, stained with DAPI (blue) and analysed by confocal microscopy for GFP fluorescence (white). Scale bar = 100 μm.

To look in more detail at HSV1 infection in the absence of nectin1, KO cells were infected with HSV1 expressing the virus protein VP22 as a GFP fusion protein (GFP-22), and analysed by flow cytometry for GFP at 8 h in comparison to infected nTERT cells. These results broadly confirmed the entry assay, indicating a massive reduction in GFP-22 positive cells in the absence of nectin1 (Fig 2B). Nonetheless, Western blotting of a high multiplicity time-course of infection further indicated that KO cells were able to support low level virus protein expression, but none of the proteins tested, including immediate-early proteins ICP4 and ICP0 were detectable much before 12 h, compared to nTERT cells where they were readily detectable by 4 h (Fig 2C). Imaging of GFP22 fluorescence in nTERT and KO cells infected in the same way revealed that these low protein levels were a consequence of HSV1 infecting a small number of KO compared to nTERT cells (Fig 2D). By contrast, high multiplicity infection of nTERT and KO cells with vaccinia virus (VACV) expressing A5-GFP revealed that VACV protein expression was unaffected by the absence of nectin1 (Fig 2E), a result that was corroborated by fluorescence microscopy of VACV infected cells where all nTERT and KO cells were shown to be A5-GFP positive (Fig 2F). Taken together, these data indicate that the majority of KO cells are refractory specifically to HSV1 infection, but that a small subpopulation retain the ability to support infection.

As final confirmation that the phenotype of these cells was a consequence of nectin1 knockout, the KO2 cells were engineered to stably express nectin1 tagged at its C-terminus with the V5 epitope. Appropriate expression of nectin1V5 was first confirmed by transient transfection of nTERT and HeLa cells followed by Western blotting for V5 (Fig S1A). This showed a V5-tagged protein which was expressed as multiple forms, and were further shown to be a consequence of *N-*linked glycosylation by treatment with the deglycosylation enzyme PNGaseF (Fig S1B). Immunofluorescence of nectin1V5 expressing nTERT cells also confirmed that V5-tagged nectin1 localised to the plasma membrane (Fig S1C). Following clonal selection of a KO2 line that had been stably transfected with the nectin1V5 plasmid, cells were tested for nectin1V5 expression by Western blotting, which indicated expression of a V5-tagged protein of the appropriate molecular weight and of similar glycosylation pattern to that seen in transiently transfected cells (Fig 3A). Flow cytometry and immunofluorescence for cell-surface nectin1 indicated that these cells expressed high levels of nectin1 on the plasma membrane in comparison to nTERT cells, albeit at levels that varied from cell to cell (Fig 3B & 3C). Imaging of permeabilised cells stained with antibodies for nectin1 and V5, which should be cell surface and cytoplasmic respectively, confirmed correct localisation of nectin1V5 (Fig 3D). Moreover, Western blotting of nectin1V5 cells infected with HSV1 expressing GFP22 indicated that in comparison to KO2 cells, these cells had regained the ability to support infection to a level similar to nTERT cells (Fig 3E), a result that was confirmed by GFP fluorescence which in contrast to KO2 cells, was present in the full population of nectinV5 cells (Fig 3F).

**Figure 3.**
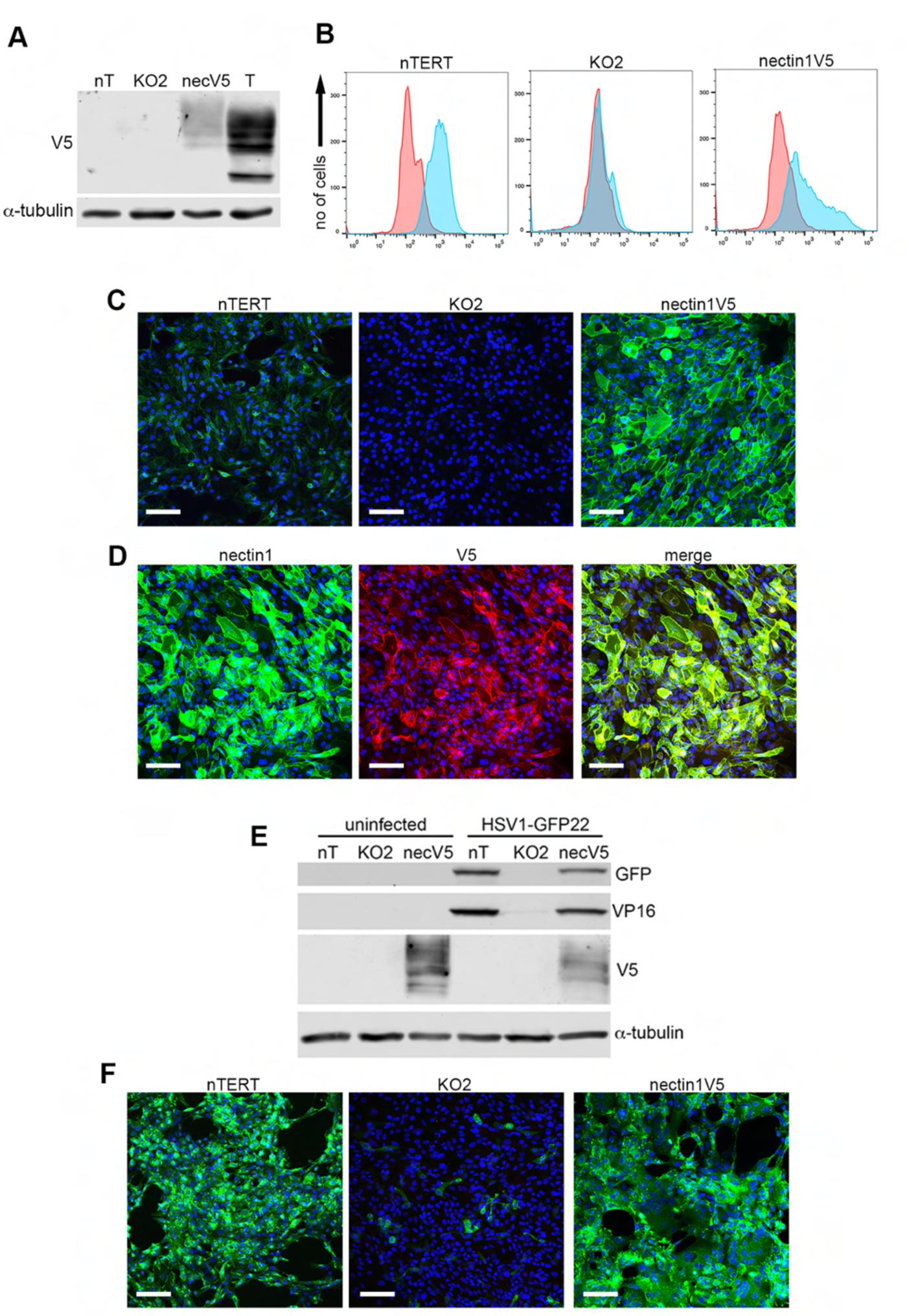
HSV1 infection is rescued in nectin1 knockout cells stably expressing nectin1V5. **(A)** KO2 cells that had been selected for stable nectin1V5 expression (necV5) were analysed by SDS-PAGE and Western blotting with antibody for V5 and *α*-tubulin alongside nTERT (nT) and KO2 cells. T, nTERT cells transiently transfected with plasmid expressing nectin1V5. **(B)** Flow cytometry of non-permeabilised nTERT, KO and nectinV5 cells was carried out using nectin1 antibody to measure nectin1V5 at the cell surface. **(C)** nTERT, KO2 and nectin-V5 cells were grown on coverslips, fixed without permeabilisation and stained with nectin1 antibody (green). **(D)** Nectin1V5 cells grown on coverslips were fixed, permeabilised and stained with antibodies to nectin1 (green) and V5 (red). Scale bar = 100 μm. **(E)** nTERT, KO and nectin1V5 cells were infected with HSV1 expressing GFP-22 MOI 5 and analysed by SDS-PAGE and Western blotting after 12 h for GFP, VP16, V5 and *α*-tubulin. **(F)** As for (E) but cells were grown on coverslips, and infected cells were fixed at 8 h nuclei stained with DAPI (blue) and cells imaged for GFP fluorescence (green). Scale bar = 100 μm.

### nTERT keratinocytes support ongoing HSV1 infection in the absence of nectin1

Closer examination of the infected cell profiles shown in Fig 2B revealed that a small number of both KO cell lines expressed high levels of GFP-22 compared to 95% of nTERT cells (Fig 4A, high GFP-22), but that an equivalent population of cells also expressed a very low level of GFP-22 (Fig 4A, low GFP-22). This result was reproduced with Wt HSV1 which were infected at high multiplicity and examined by immunofluorescence for the immediate-early protein ICP4. ICP4 exhibits a transition in localisation according to the time of infection: initially in a diffuse nuclear pattern, followed by localisation to nuclear replication compartments, and finally at later times in a cytoplasmic punctate pattern; it can therefore be used to monitor the kinetic stage of HSV1 infection. In this case, at 4 h, the majority of nTERT cells but only a subpopulation of KO cells expressed ICP4 (Fig 4B, compare nTERT and KO), whereas by 16 h, the number of ICP4 positive KO cells in the monolayer had greatly increased (Fig 4B, KO). A time course of nTERT and KO cells infected at high multiplicity with HSV1 expressing GFP-22 and stained for ICP4 revealed that the majority of nTERT cells expressed ICP4 by 4 h and GFP-22 by 6 h (Fig 4C), in line with the known replication kinetics of HSV1 in nTERT keratinocytes [5]. By contrast, only a small number of KO cells expressed ICP4 at 4 h and a similar number expressed GFP-22 by 8 h (Fig 4C, KO). Nonetheless, by 8 h, many more cells were positive for ICP4 in its early diffuse nuclear localisation pattern, without expressing GFP-22, suggesting an early infection (Fig 4C, KO 8 h). Taking together the flow cytometry and microscopy results, this suggests that in the absence of nectin1, a small number of nTERT cells are infected with HSV1 with similar kinetics to nectin1-expressing nTERT cells, while a second population exhibits a delayed infection.

**Figure 4.**
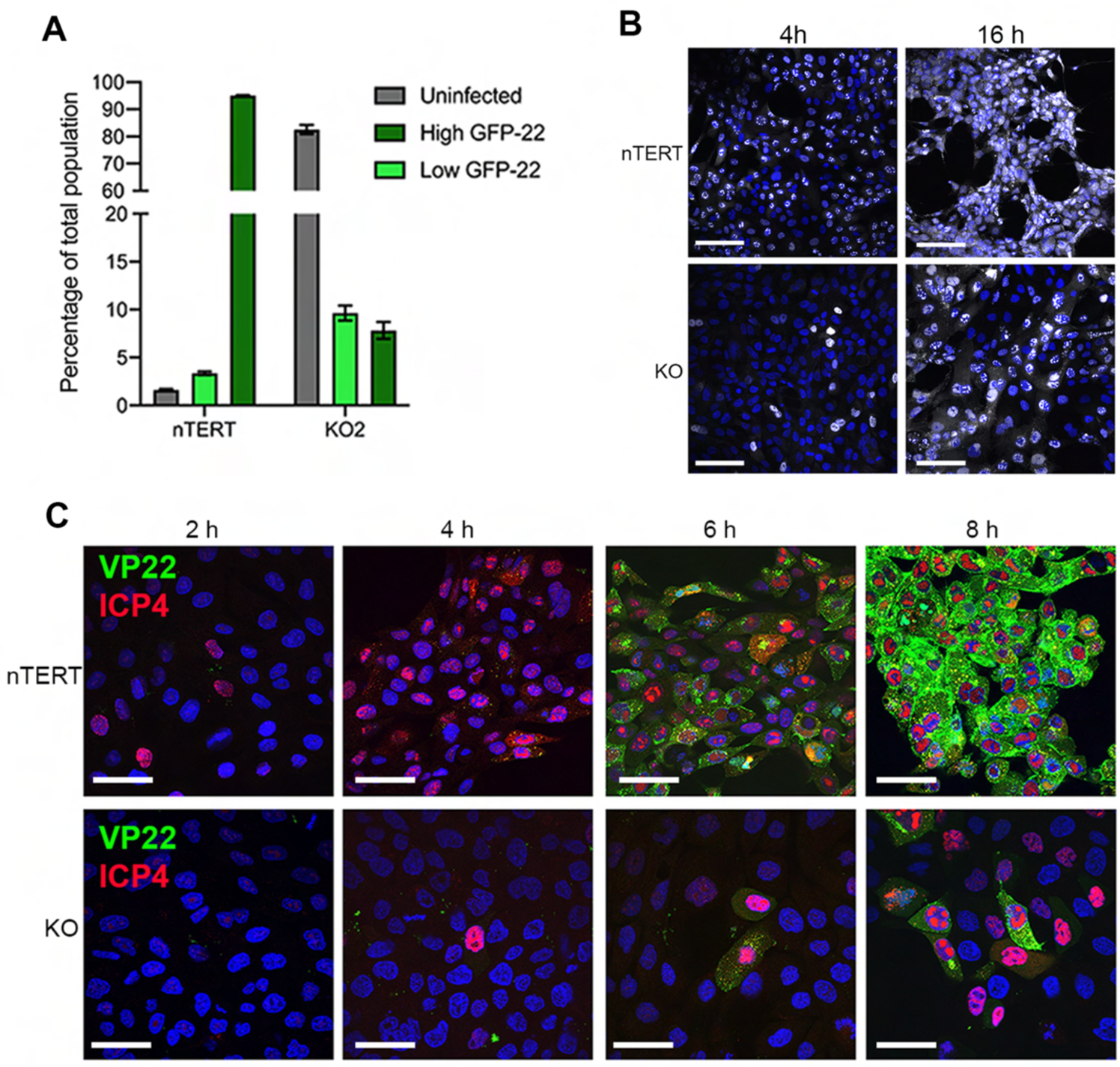
HSV1 exhibits delayed infection in nectin1 KO cells. **(A)** GFP positive cells were quantitated by flow cytometry and separated into high or low GFP expression in nTERT or KO2 cells infected with HSV1 GFP-22. **(B)** nTERT and KO cells grown on coverslips were infected with HSV1 Sc16 at MOI 5 and fixed at 4 h or 16 h. Cells were permeabilised and stained for the immediate-early protein ICP4 (white) and nuclei stained with DAPI (blue). Scale bar = 100 μm. **(C)** nTERT and KO cells grown on coverslips were infected with HSV1 GFP-22 (green) at MOI 5 and fixed at the indicated times. Cells were permeabilised and stained for ICP4 (red) and nuclei were stained with DAPI (blue). Scale bar = 50 μm.

This pattern of high and low-level infected cells in nectin1 KO cells could be explained by delayed infection from the initial virus inoculum or the spread of newly replicated virus from the initially infected cells into a larger population of uninfected cells. To determine if a new round of genome replication was required, nTERT and KO cells were infected with Sc16 in the absence or presence of DNA replication inhibitor AraC, fixed at 16 h and stained for ICP4, indicating that AraC blocked the increase in the number of infected cells (Fig 5A, compare -AraC and + AraC). Moreover, prolonged exposure of KO cells to the HSV1 inoculum in the presence of AraC had no effect on the number of initially infected cells, indicating that the failure to infect a larger proportion of cells in the first instance was not simply a consequence of a reduced rate of entry (Fig 5A, KO +AraC, compare 1h and 16h). To confirm that newly produced virus was required for the observed delay in infection, we made use of a virus expressing GFP in place of the nuclear egress protein UL34 (*Δ*UL34). This deletion mutant expresses all proteins including late proteins as normal, and capsids assemble in the nucleus, but they fail to translocate to the cytoplasm for subsequent envelopment [20]. In this case, while all nTERT cells expressed both ICP4 and GFP at this late time when infected at high multiplicity (Fig 5B, nTERT), only the original susceptible subpopulation of KO cells expressed these proteins (Fig 5B, KO), indicating that ICP4 expression in the later population of cells required the production of infectious virions.

**Figure 5.**
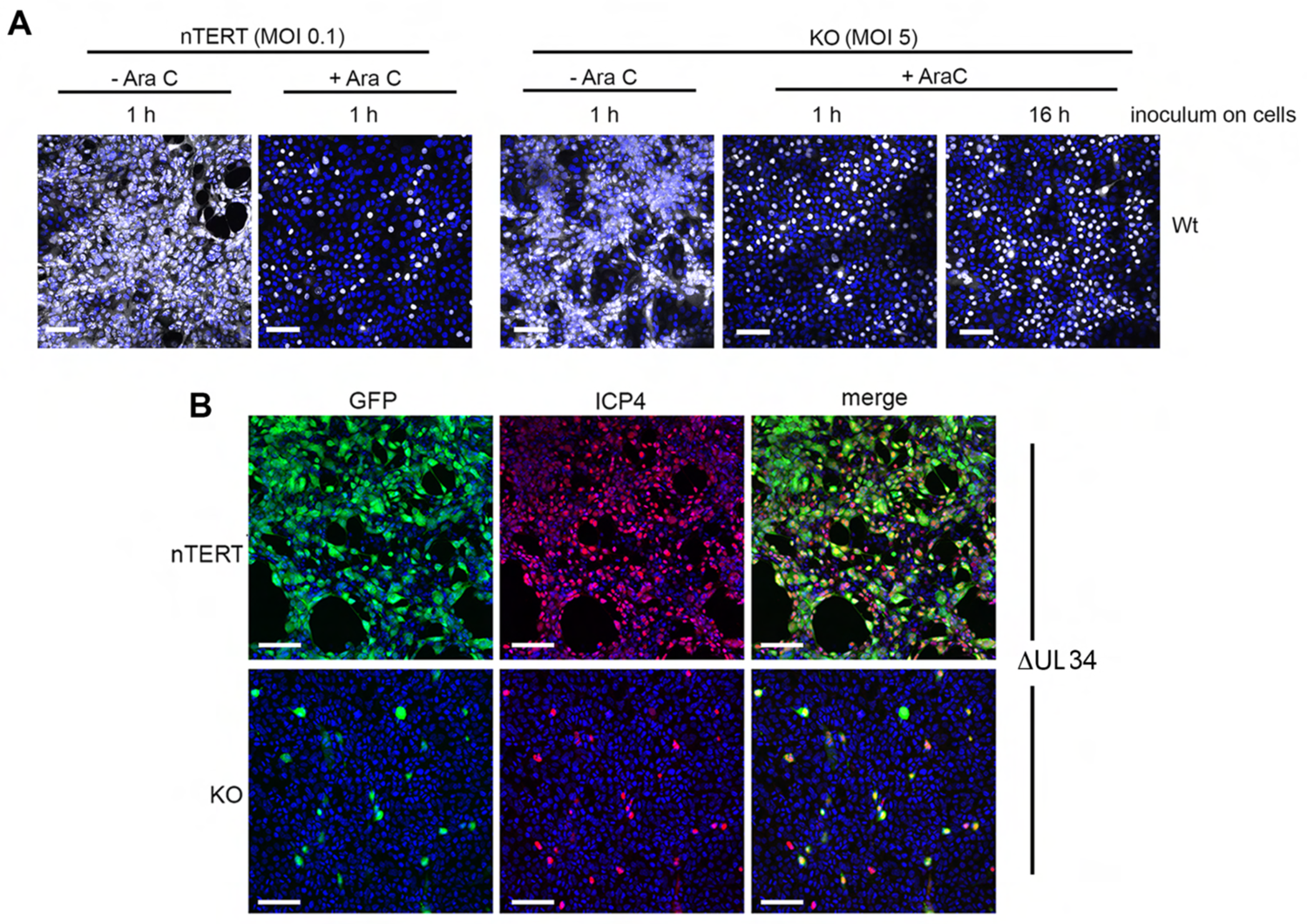
Delayed HSV1 infection of nectin1 KO cells requires virus replication. **(A)** nTERT and KO cells were infected with HSV1 Sc16 at MOI 0.1 or 5 respectively in the absence or presence of 100 ng/ml AraC. Inoculum was removed after one hour as normal (1 h), or left on for the course of the infection (16 h). Cells were fixed at 16 h and stained for ICP4 (white) and nuclei were stained with DAPI (blue). Scale bar = 100 μm. **(B)** nTERT and KO cells grown on coverslips were infected with HSV1 *Δ*UL34 at MOI 5 and fixed at 20 h. Cells were permeabilised and stained for ICP4 (red) and nuclei were stained with DAPI (blue). GFP expressed in place of UL34 is in green. Scale bar = 100 μm.

### HSV1 spreads to neighbouring nectin1 KO cells

The above results indicate that virus infectivity is able to spread through nectin1 KO cells. This was unexpected because the initial susceptibility of these cells was around 5%. To assess how this new round of infection was transmitted through the monolayer, conditions for detecting virus spread through the cells were first established. In this case, low multiplicity infections of nTERT cells in the absence and presence of AraC were fixed at different times after infection and stained for ICP4 by immunofluorescence (Fig S2). This indicated that under these conditions spread was detectable at 12 hours, and that as expected from the fully susceptible nature of these cells, this spread occurred in the immediate locality of the initially infected cell, spreading in to all cells over 16 h (Fig S2). Using the same conditions on KO cells, we were also able to detect local infection around the initial infected cell which was absent in AraC treated cells (Fig 6A). Moreover, high magnification imaging of KO cells infected with HSV1 expressing GFP22 and stained for ICP4 revealed that cells containing a low level of ICP4 in the nucleus were frequently located beside cells expressing high level of ICP4 and GFP-22 (Fig 6B, arrowed). Additional imaging of cells infected with HSV1 expressing GFP-26 also allowed the visualisation of virus capsid localisation in ICP4-positive nectin1 KO cells (Fig 6C). In both nTERT and KO cells, individual progeny capsids were clearly detectable not only in the initially infected cells, but also in cells surrounding the first infected cell, some of which were already producing ICP4 (Fig 6C, arrowed). These results confirm that while around 95% of the nectin1 KO cells are resistant to HSV1 infection from virions in the medium, they are fully susceptible to infection from virus transmitting from adjacent cells.

**Figure 6.**
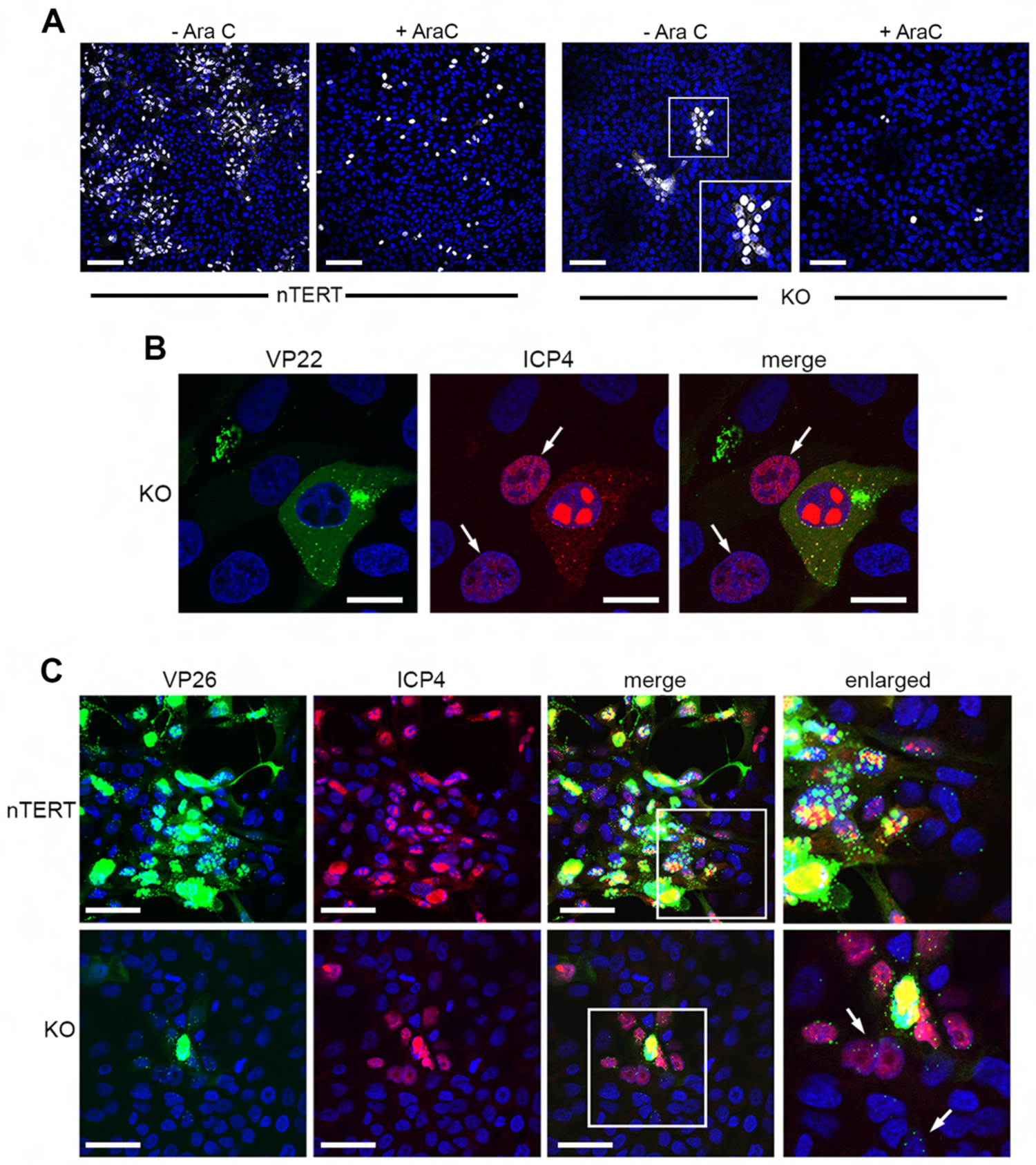
HSV1 infection spreads in nectin 1 KO cells. **(A)** nTert and KO cells were infected with HSV1 Sc16 at MOI 0.05 in the absence or presence of 100 ng/ml AraC to block virus genome replication. Cells were fixed at 12 h and stained for ICP4 (white) and nuclei were stained with DAPI (blue). Scale bar = 100 μm. **(B)** nTERT and KO cells grown on coverslips were infected with HSV1 GFP-22 (green) at MOI 5 and fixed at 8 h. Cells were permeabilised and stained for ICP4 (red) and nuclei were stained with DAPI (blue). Scale bar = 20 μm. White arrows indicate cells with early expression of ICP4. **(C)** nTERT and KO cells grown on coverslips were infected with HSV1 GFP-VP26 at MOI 5 and fixed at 12 h. Cells were permeabilised and stained for ICP4 (red) and nuclei were stained with DAPI (blue). GFP-VP26 is in green. Scale bar = 50 μm. Arrow points to cells with GFP-tagged capsids next to the original infected cell.

### Phenotype of HSV1 lacking glycoproteins gE or gI in the absence of nectin1

HSV1 has two modes of spreading between cells: either by extracellular release and spread through the liquid medium to re-enter uninfected cells, or via direct cell-to-cell spread from already infected cells [12]. Infection by cell-to-cell transfer is known to involve glycoproteins gE and gI, which are both non-essential in tissue culture but essential *in vivo* [14, 15], and viruses lacking either of these glycoproteins form small plaques [15]. Hence, we next tested the ability of HSV1 mutants in gE or gI to infect and spread in the nectin1 KO cells. In this case we used two different gI mutant viruses, both in a Sc16 background: one called *Δ*gI that was previously constructed by insertion of the *β*-galactosidase gene into the gI-encoding Us7 open reading frame [15] (Fig S3A) and one called *Δ*gIGFP that has been constructed by ourselves, in which Us7 has been replaced by the GFP open reading frame (Fig S3B & S3C). nTERT and KO cells were infected with each of these viruses in the absence or presence of AraC to determine their ability to infect the initial susceptible population of KO cells, and to establish they can spread into neighbouring cells in a similar fashion as the Wt virus. These viruses were all able to infect nTERT and KO cells with the same efficiency as Wt, confirming that the absence of gE or gI had little effect on infection from without in the presence or absence of nectin1 (Fig 7, + AraC). Infection in the absence of AraC indicated that the glycoprotein mutant viruses were all able to spread in nTERT, although *Δ*gE appeared to spread less efficiently (Fig 7, nTERT). Likewise, the *Δ*gE and *Δ*gIGFP viruses spread into surrounding KO cells, but at a reduced rate compared to Wt virus (Fig 7, KO). By contrast, the *Δ*gI virus failed to spread into surrounding cells in the time frame of this experiment suggesting its phenotype was different to to that of *Δ*gIGFP virus (Fig 7, KO).

**Figure 7.**
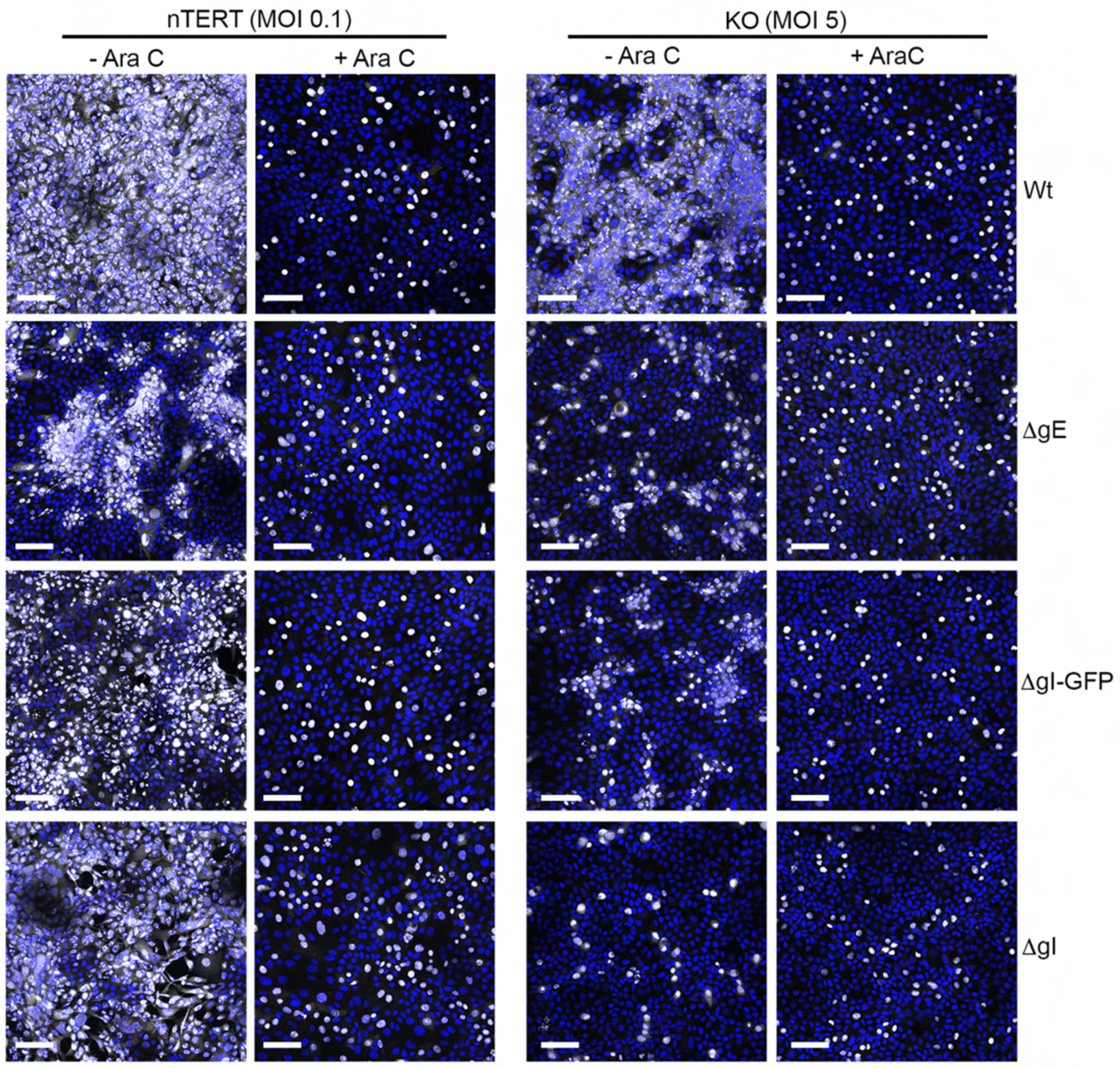
Spread phenotype of HSV1 lacking glycoprotein E or glycoprotein I in nectin1 KO cells. nTERT and KO cells grown on coverslips were infected with HSV1 viruses as indicated at MOI 0.1 for nTERT and MOI 5 for KO cells in the absence or presence of 100 ng/ml AraC. Cells were fixed at 16 h, permeabilised and stained for ICP4 (white). Nuclei were stained with DAPI (blue). Scale bar = 100 μm.

These contradictory results for the *Δ*gI viruses led us to investigate the composition of virions made by both of these viruses. Wt, *Δ*gI and *Δ*gIGFP virions were purified from infected HaCaT cells and analysed by SDS-PAGE followed by Coomassie blue staining to equalise loading of virions and Western blotting (Fig S4A & B). Strikingly, we found that *Δ*gI virions contained a very low level of gD compared to Wt virions, while *Δ*gIGFP virions contained roughly normal levels of gD (Fig S4B). Moreover, Western blotting of infected cell lysates indicated that the low level of gD in the *Δ*gI virions correlated with a low level of gD expression (Fig S4C). This suggests that the *Δ*gI virus is unable to spread in KO cells not because of an absence of gI but because of its greatly reduced gD expression, and by extension then that gD is required for spread in nectin1 KO cells.

Extended spread of Wt, *Δ*gE and *Δ*gIGFP viruses was next determined by infecting nTERT and KO monolayers with a low number of virus plaque forming units, followed by fixing and staining for ICP4 at 24 h and 48 h (Fig 8). Wt foci were larger in nTERT cells compared to KO cells, indicating that although able to transmit, this process occurred more slowly in the absence of nectin1 (Fig 8A & 8C). *Δ*gE and *Δ*gIGFP viruses were able to spread in nTERT cells but more slowly than Wt, as would be expected from their known phenotypes (Fig 8A). Likewise, these viruses were able to spread over time in KO cells, but this was also at a slower rate than Wt virus (Fig 8C). When examining these foci, it was clear that the *Δ*gE and *Δ*gIGFP viruses were spreading in nTERT cells to form characteristic “comet tails” [12] a feature caused by virus moving through the extracellular medium (Fig 8A). On the other hand, Wt virus appeared to spread predominantly in the locality of the original infected cell, with only limited evidence of extracellular movement to infect distant cells and cause comet tails (Fig 8B). In KO cells, none of these viruses appeared to spread to cause comet tail formation (Fig 8C).

**Figure 8.**
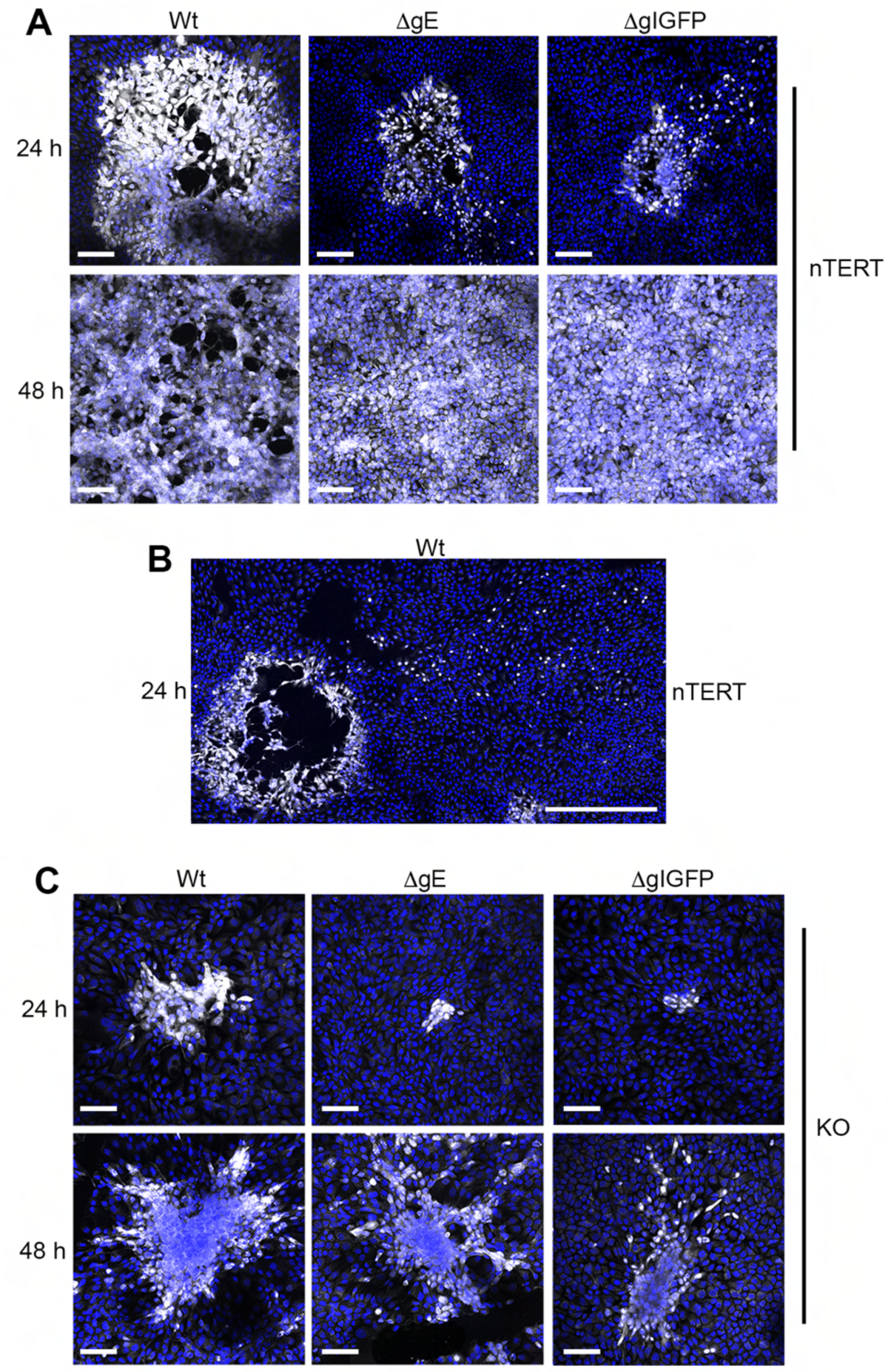
nTERT **(A & B)** and KO **(C)** cells grown on coverslips were infected with 50 or 2000 plaque forming units respectively of HSV1 Wt (Sc16), *Δ*gE or *Δ*gIGFP, fixed at 24 h or 48 h and stained for ICP4 (white). Nuclei were stained with DAPI (blue). Scale bar = 100 μm (A & C) or 500 μm (B).

### Direct cell-to-cell transmission of HSV1 occurs in the absence of nectin1

To confirm if HSV1 spreads cell-to-cell in nectin1 KO cells, low multiplicity infections of HSV1 strain Sc16, *Δ*gE, *Δ*gIGFP together with *Δ*gI were carried out in nTERT and KO cells in the presence of human serum, used here to neutralize extracellularly released virus, thereby eliminating extracellular spread and allowing virus spread only by direct transfer between cells in contact [12]. As in the absence of human serum, the glycoprotein deletion viruses were unable to spread to the same extent as Wt virus, but *Δ*gE, *Δ*gIGFP and *Δ*gI all spread similarly in nTERT cells (Fig 9A). In KO cells, this differential spread was similar for *Δ*gE and *Δ*gIGFP, but as seen above in Fig 7, the *Δ*gI virus all but failed to spread into surrounding cells (Fig 9A). Measurement of these infected cell foci confirmed that the inhibition of extracellular spread by human serum had no effect of the spread area of Wt infection in either nTERT or KO cells (Fig 9B), while the absence of gE or gI causes a similar relative reduction on foci size in nTERT and KO cells (Fig 9C). This confirms that HSV1 maintains the ability to spread in keratinocytes in the absence of nectin1, but that this spread occurs in a gE-gI-dependent fashion by direct cell-to-cell transmission rather than extracellular release and local reinfection. Moreover, given that the *Δ*gI virus expresses very low levels of gD and cannot spread in the absence of nectin1, the nectin1-independent spread route is highly dependent on gD.

**Figure 9.**
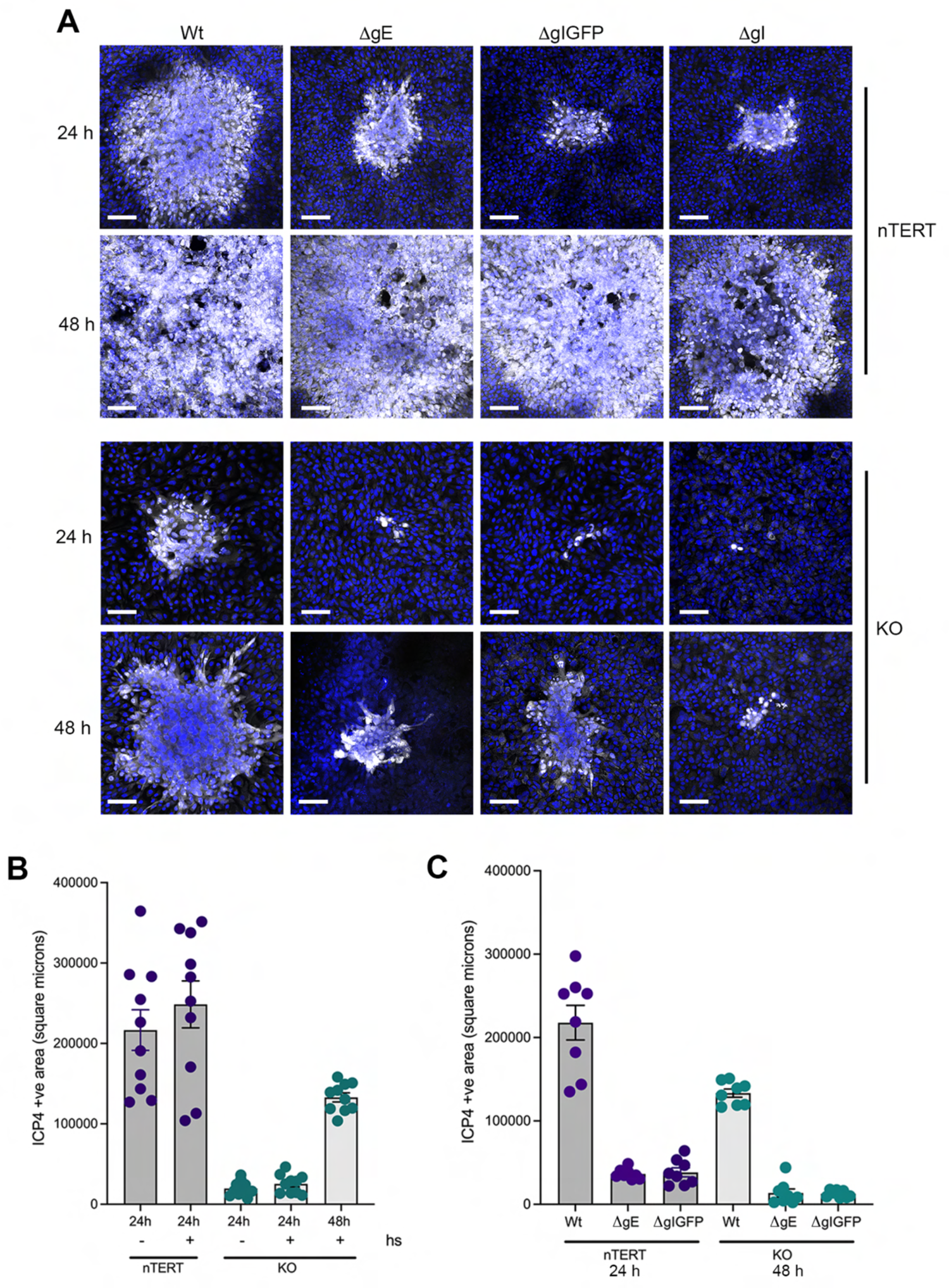
HSV1 spreads via cell-to-cell transmission in the absence of nectin1. **(A)** nTERT and KO cells grown on coverslips were infected with 50 or 2000 plaque forming units respectively of HSV1 viruses as indicated in the presence of 10% human serum to neutralise extracellular virus and block extracellular spread. Cells were fixed at the indicated times, stained for ICP4 (white) and nuclei stained with DAPI (blue). Scale bar = 100 μm. **(B)** The area of ten ICP4-positive Wt foci on nTERT and KO cells infected in the absence or presence of human serum (hs) was measured using NIH ImageJ**. (C)** The area of eight ICP4-positive foci of Wt, *Δ*gE and *Δ*gIGFP viruses on nTERT and KO cells infected in the presence of human serum (hs) was measured using NIH ImageJ.

Although our previous data had suggested that HVEM has little role in entry into nTERT keratinocytes when nectin1 is present [5], the above results led us to investigate if depletion of HVEM in the nectin1 KO cells affected virus entry and/or spread in the absence of nectin1. Additionally, we wanted to determine if the previously identified spread factor PTP1B [17] played a role in the nectin1-independent spread of HSV1 in keratinocytes. RT-qPCR of HVEM mRNA indicated that HVEM and PTP1B transcripts are expressed at similar levels in both nTERT and KO cells (Fig S5A & S5B). nTERT and KO cells were transfected with control, HVEM or PTP1B siRNAs, and RT-qPCR was used to confirm efficient depletion of each transcript (Fig S5C). Subsequent infection with Wt virus, followed by fixation and staining for ICP4 at 8 h and 16 h showed that neither initial entry nor spread of HSV1 in nTERT or KO cells was altered in cells depleted of HVEM or PTP1B (Fig 10), and therefore the nectin1-independent spread pathway that functions in KO cells does not appear to involve either of these previously identified factors.

**Figure 10.**
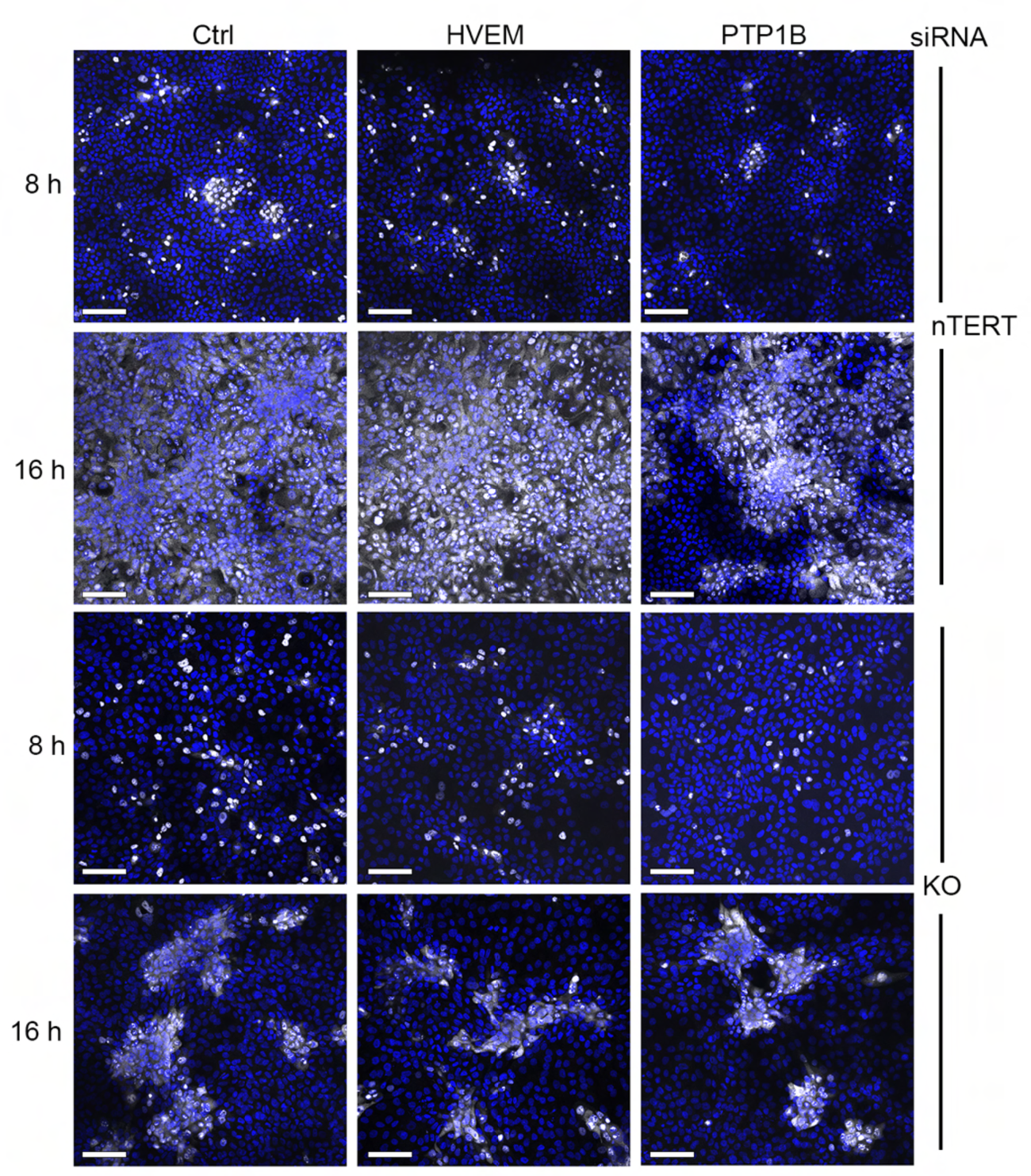
HSV1 spread in the absence of nectin1 does not require HVEM or PTP1B. nTERT and nectin1 KO cells were transfected with control (Ctrl), HVEM or PTP1B siRNA. After 48 h, nTERT and KO cells were infected with HSV1 Sc16 MOI 0.1 or 5 respectively, fixed at 8 or 16 h and stained for ICP4 (white). Nuclei were stained with DAPI (blue). Scale bar = 100 μm.

In line with the above results, titration of HSV1 on nTERT and nectin1 KO cells indicated that HSV1 was able to form plaques on nectin1 KO cells, although as expected from the foci staining experiments, these were smaller than plaques on nTERT cells (Fig 11A & 11B, HSV1). Moreover, the relative titre of HSV1 was around 50-fold lower in KO cells compared to nTERT cells (Fig 11C), reflecting the relative efficiency of initial infection seen above by flow cytometry and confocal microscopy. Interestingly, HSV2 behaved in the same way as HSV1 in all these assays, (Fig 11A to 11C, HSV2), while immunofluorescence of HSV2-infected monolayers confirmed the similarity between the two HSV types in the absence of nectin1 (Fig 11D). In all, this confirms that in spite of only being able to infect a small subpopulation of cells from the extracellular medium in the absence of nectin1, once in those cells, both HSV1 and HSV2 can go on to spread into the surrounding nectin1 KO cells which had been initially resistant to HSV1 entry.

**Figure 11.**
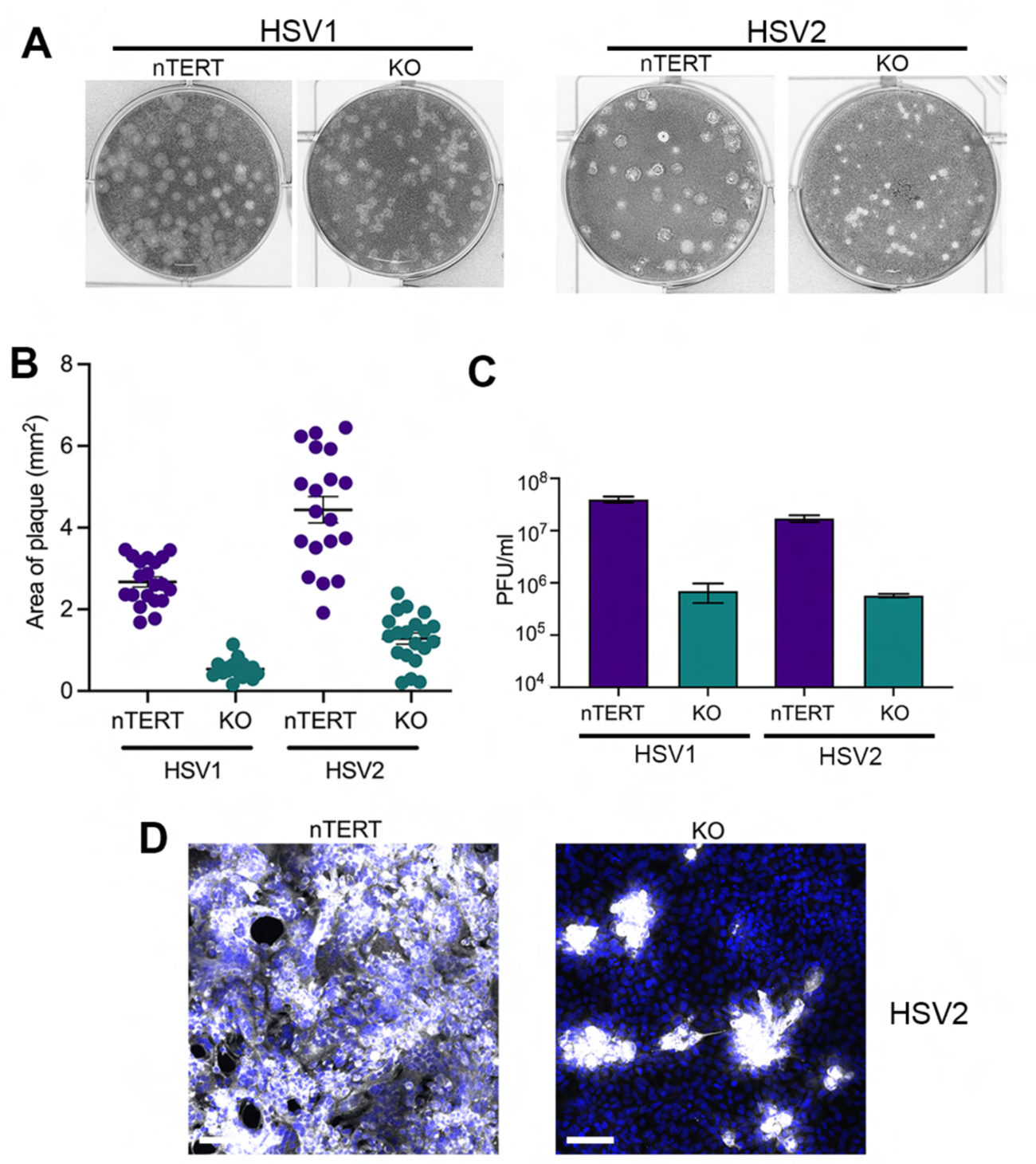
HSV types 1 and 2 form plaques on nectin1 KO cells. **(A)** HSV1 strain Sc16 and HSV2 strain HG52 was titrated onto nTERT and KO cells in the presence of 3% CMC, and fixed and stained with crystal violet 3 days later**. (B)** The area of ten HSV1 and HSV2 plaques on nTERT and KO cells shown in (A) was measured using NIH ImageJ**. (C)** The relative titre of HSV1 and HSV2 on nTERT and KO cells (mean±SEM, *n* = 3). **(D)** nTERT and KO cells were infected with HSV2 MOI 5 and fixed at 16 h. Cells were permeabilised and stained with an antibody to glycoprotein D (green) and nuclei stained with DAPI (blue). Scale bar = 100 μm.

## Discussion

Like many viruses that infect host epithelia, HSV has two possible transmission routes: entry of cell-free particles from the extracellular environment, or direct movement between cells [13, 14]. Despite the second route being important for spread in the host, the cell biology and molecular players involved in this mechanism are poorly understood. Here, we have not only proved conclusively that nectin1 is the major receptor for HSV1 to gain access from the extracellular environment in to human keratinocytes – the main cell type in the epithelia that HSV1 infects - but we have also demonstrated that cell-to-cell spread of HSV1 in these cells occurs in the absence of this receptor. Moreover, as anticipated from the original seminal report on the discovery of nectin1 as a prime candidate for an HSV entry receptor [7], HSV2 has the same entry requirements and spread properties as HSV1 in human keratinocytes. Consequently, once HSV has gained entry in to the small population of susceptible nectin1 KO cells, it is able to spread unrestrained and form plaques by moving directly from cell-to-cell, not by extracellular release. This unexpected result shows for the first time that despite being the major receptor for entry from the extracellular environment, nectin1 is not required for HSV to move between keratinocyte cells in culture. Additionally, the second major HSV receptor known as HVEM [8] was also shown to be dispensable for cell-to-cell spread in the absence of nectin1 suggesting that this route of transmission occurs via a novel pathway which despite relying on gD does not require either of the two main gD receptors.

There are two possible means of intact virus particles undergoing cell-to-cell transfer. The first mechanism involves retention of the virion at the plasma membrane after egress from the infected cell, followed by binding to and rapid entry into the uninfected cell, similar to that seen in the HIV-induced T-cell virological synapse [21] or the projection of vaccinia virus particles into uninfected cells on actin tails generated from inside the infected cells [22]. Multiple ultrastructural studies have indicated that vast numbers of HSV1 virions line up between the plasma membranes of adjacent cells after egress from a range of cell types, indicating that external virions maintain interactions with the cell surface, providing a concentrated source of particles for potentially rapid transfer into uninfected cells. The second mechanism which we cannot exclude, is the transfer of virus infectivity between either pre-existing or virus-induced intercellular connections, formed between nectin1 KO cells. As such, this movement of infectivity could involve either fully assembled virions or virion sub-particles, for example in the form of unenveloped capsids moving between cell-to-cell contacts. Although results with the *Δ*UL34 virus confirmed that capsid export from the nucleus to the cytoplasm was required for the spread of infection, and imaging of GFP-tagged capsids showed capsid movement from infected into uninfected nectin1 KO cells, it is still to be formally proven that infectivity in these cells is transmitted in an enveloped virion rather than a naked capsid. Examples of such sub-particle transmission already exist in plant viruses, where genomes of, for example, tobacco mosaic virus are transferred between cells through plasmodesmata which have been dilated by the action of the virus movement protein [23]. Similar mechanisms of spread through intercellular connections known as tunnelling nanotubes have also been implicated in the spread of respiratory viruses including measles virus (MeV) and respiratory syncytial virus, which infect the simple columnar epithelia of the human respiratory tract [24, 25]. In addition, MeV has also been shown to spread by cell contact in neurons [26]. Interestingly, there is recent evidence that the animal alphaherpesvirus bovine herpesvirus type1 may transmit through nanotubes in culture [27].

Given the known role of the non-essential glycoproteins gE and gI in cell-to-cell transfer in epithelial and neuronal cells [14, 28–30], it is maybe not surprising that the nectin1 independent cell-to-cell spread mechanism of HSV1 was shown here to involve these glycoproteins. Although the functions of gI and gE have not been fully defined, they are essential *in vivo* [14, 15], and are therefore candidate factors to facilitate the transfer of virions from one keratinocyte to another. Indeed, they have previously been shown to be involved in the sorting of nascent virions to cell junctions in polarised epithelial cells [31]. In addition, they are required for anterograde transport of virions in neuronal axons [32, 33], by recruiting specific kinesins for microtubule movement [34, 35], a mechanism that could also function in epithelial cells. However, *Δ*gI and *Δ*gE virions are efficiently released from epithelial cells in culture suggesting that transport to the plasma membrane is unhindered in these cells [14, 28], and it is therefore unlikely that this kinesin-mediated microtubule transport role is generally applicable for delivery to the plasma membrane of epithelial cells. Nonetheless, this mechanism may channel virions for delivery on microtubule tracks to the specific areas of the plasma membrane that form cell-to-cell contacts, thereby potentially delivering them to the correct site for rapid spread. Moreover, it was recently suggested that inhibition of the ER-bound tyrosine phosphatase, PTP1B, specifically blocked cell-to-cell spread of HSV1, suggesting that tyrosine phosphorylation regulates either the cellular or viral trafficking proteins that make up the cell-to-cell spread machinery [17]. However, those experiments were carried out in HaCaT cells, and in our hands PTP1B was not required for cell-to-cell spread as shown by siRNA depletion here and use of the PTP1B inhibitor (data not shown). Future live cell and ultrastructural studies of the nectin1 KO keratinocytes will help to delineate how these different features integrate to facilitate cell-to-cell spread of HSV1 and how the proteins of the virus spread machinery including gI and gE, contribute to this process.

Cell-to-cell spread of virus in the host allows the virus to avoid neutralizing antibodies and some intrinsic/innate antiviral responses whilst providing an efficient and rapid way for small numbers of virus particles or sub-particles to disseminate infection. It is of note then that clinical isolates of HSV1 have the propensity to be cell-associated when first isolated, whereas lab strains often produce large amounts of free virus [12]. This is similar to the situation in human cytomegalovirus, a betaherpesvirus that is predominantly cell-associated in the host, but rapidly loses a large part of its genome over as few as three passages in cultured fibroblasts, resulting in virus that is less cell-associated and reduced in pathogenicity [36]. Moreover, varicella zoster virus (VZV), another human alphaherpesvirus which infects keratinocytes in the host to cause chickenpox or shingles, is highly cell-associated in culture [37], and unlike other alphaherpesviruses which can spread by the extracellular route, VZV does not have a homologue of HSV1 gD for receptor binding from the outside [37]. Taken together, this indicates that the cell-to-cell spread machinery is vital for VZV infection and as such, it is intriguing to speculate that our nectin1 KO keratinocytes may recapitulate the natural spread mechanism of VZV but in an HSV background.

There is growing evidence that human keratinocytes express a much higher level of nectin1 than many other human cell-types used for HSV1 studies [10]. Nonetheless, although the vast majority of nectin1 KO cells were found here to be resistant to HSV1 entry from outside, a small proportion remained susceptible, indicating entry via another route. The number of cells infected could reflect a requirement for the cells to be, for example at a certain stage of the cell-cycle for virus entry to occur in the absence of nectin1. However, the fact that prolonged exposure to the virus inoculum did not result in an increased number of initially infected cells suggests this is unlikely to be the case. Alternatively, another receptor may be expressed at the cell surface in this small population of cells allowing virus binding and entry into those cells. Nonetheless, depletion of the alternative gD receptor HVEM in the nectin1 KO cells had little effect on virus entry in to the susceptible KO cells, and therefore other candidate receptors such as nectin2 [38], or the previously identified gB receptor paired immunoglobulin-like type 2 receptor PILR*α* [39] will need to be investigated. Whatever the mechanism and receptor involved, it is clear that these pathways of entry are minor components of the entry process, and that the expression of nectin1 at the keratinocyte cell surface facilitates the efficient entry of HSV types 1 and 2 from the extracellular environment.

The results presented here imply that HSV1 requires nectin1 for host-to-host transmission, facilitating virus entry into the host epithelium. However, there is a real possibility that once the virus has gained access into its first target cell, nectin1 is not required in virus spread at least within the epithelium. It is noteworthy that HSV1 infects multi-layered, stratified epithelia, where individual keratinocytes form cell-to-cell contacts in all dimensions. Work is now needed on three-dimensional raft cultures of keratinocytes to determine if our results in keratinocyte monolayers translate to a model tissue and ultimately the host. Our nectin1 KO cells – and in particular the ease with which we can image GFP-tagged capsid movement in these cells – provide new scope to study the spread of HSV types 1 and 2 in all dimensions in the absence of confounding effects from released extracellular virus.

## Materials and Methods

### Cells and Viruses

nTERT cells were cultured in 3:1 Dulbecco’s modified Eagle media (DMEM) to Hams F12 media supplemented with RM+ supplement (10 ng/ml mouse epidermal growth factor (Serotec), 1 ng/ml cholera toxin (Sigma), 400 ng/ml hydrocortisone (Sigma), 5 μg/ml apo-transferrin (Sigma) and 13 ng/ml liothyronine (Sigma), 50 U/ml penicillin streptomycin and 10% foetal bovine serum (FBS). Vero, HeLa and HaCaT cells were cultured in DMEM supplemented with 50 U/mL penicillin/streptomycin and 10% FBS. All HSV viruses except *Δ*UL34 were routinely propagated in Vero cells in DMEM supplemented with 50 U/mL penicillin/streptomycin and 2% FBS. Plaque assays were carried out in nTERT in RM+ media supplemented with 2% FBS, 50 U/mL penicillin/streptomycin with 3% carboxymethyl cellulose (CMC) as an overlay. Cell-to-cell spread assays were carried out using pooled human serum (BioIVT) at a concentration of 10% to neutralise extracellular virus. Arabinofuranosyl Cytidine (AraC) was used at a concentration of 100 ng/ml in media to inhibit HSV1 genome replication. VACV was propagated in RK13 cells and titrated on BSc1 cells.

HSV1 strain Sc16 was routinely used in this study [40]. Strain s17 expressing GFP-VP22 or GFP-VP26, and Sc16110lacZ have been described previously [18, 41, 42]. *Δ*gI and *Δ*gE viruses have been described before [15] Sc16 expressing GFP in place of Us7 (*Δ*gIGFP) was constructed using homologous recombination (Fig S5). The HG52 strain of HSV2 was used here. VACV western reserve (WR) strain expressing A5-GFP was a gift from Carlos Maluquer de Motes (University of Surrey) and has been described before [43]. HSV1 deleted for UL34 and UL34-expressing Vero cells for its propagation [20] were a gift from Richard Roller (University of Iowa). Extracellular virions were prepared as described previously [44].

### Antibodies used in this study

The following primary antibodies were kindly provided by: mouse anti-gD (LP14) and mouse anti-VP16 (LP1) Colin Crump (University of Cambridge); mouse anti-gI (3104) David Johnson (Oregon health and Science University); mouse anti-F13, Carlos Maluquer de Motes (University of Surrey). Commercially available antibodies used include: mouse anti-ICP0 (11060; Santa Cruz); mouse anti-ICP4 (Santa Cruz); mouse anti-nectin1 (R1.302; Biolegend); mouse anti-*α* tubulin (Sigma).

### SDS-PAGE and Western blotting

Samples were separated on SDS-PAGE gels from 10-14% as appropriate and transferred to nitrocellulose membranes before incubation with primary antibody. Goat anti-mouse IRDye 680RD and goat anti-rabbit IRDye 800CW (LI-COR Biosciences) secondary antibodies were used as appropriate, before blots were either imaged using an Odyssey CLx imaging system (LI-COR Biosciences) or developed using SuperSignal West Pico chemiluminescent substrate and exposed to X-ray film.

### Flow cytometry

Cell surface staining for nectin1 was performed on live cells that had been blocked by incubation in PBS with 5% FBS and 1 mM EDTA for 10 mins. Mouse anti-nectin1 was added in PBS with 5% FBS and 1 mM EDTA for 30 min. After extensive washing with PBS, Alexafluor 488 anti-mouse secondary antibody was added in PBS with 5% FBS and 1 mM EDTA and incubated for a further 30 min. Cells were then washed before being fixed in 1% PFA for 20 mins, suspended in PBS with 1% FBS and 1 mM EDTA and analysed using a BD FACS Celesta flow cytometer. Infected cells were trypsinised and fixed as above then directly analysed for GFP fluorescence. Levels of GFP-22 fluorescence were ascertained by comparison to mock and the maximum WT infected GFP levels using FlowJo software.

### CRISPR-Cas9 knockout of nectin1

Two guide RNA sequences targeting nectin-1 (GACTCCATGTATGGCTTCATCGG & GAGTCGTTCACCTGGACCACCTGG) were inserted as annealed oligonucleotides with BBS1 overhangs into BBS1 digested plasmid pX459 which expresses Cas9. Plasmids were transfected into nTERT cells in a 1:1 ratio with Lipofectamine 2000 (Invitrogen). For KO cells media was replaced 24 hours later with media containing 500 ng/μl puromycin and incubated for a further 72 h. Surviving cells were trypsinised to a single cell suspension, added to 96 well plates at a concentration of 5 cells/ml and grown as clonal populations. For KO2 cells, cells were trypsinised to a single cell suspension 24 h after transfection. Live cells were blocked and stained as for flow cytometry above, and fluorescence associated cell sorting (FACS) was performed using a FACS ARIA fusion (BD Biosciences). Nectin1 negative cells were identified by gating using FACS DIVA v8 software and single cell sorted into 96 well plates containing filtered conditioned media. Once confluent potential clones were expanded and screened for nectin1 deletion.

### Construction of keratinocytes expressing epitope-tagged nectin1

Plasmid pNectinV5 [45] was transfected into KO2 cells with Lipofectamine 2000. Seventy two hours post transfection cells were moved to a 33°C incubator for 5 days to promote homologous recombination. Live cells were blocked and nectin1 stained for flow cytometry as above. Nectin1 positive cells were then sorted into 96 well plates containing filtered conditioned media. Once confluent, potential clones were screened for V5 expression.

### Genomic sequencing

Genomic DNA was extracted from cells using the DNeasy blood and tissue kit (Qiagen). Nectin1 deletion was confirmed by PCR amplification of the 5’ region of the *NECTIN1* gene (Forward and reverse primers: GACTCAGCTGCGAGGGAGAAG & GAGCTGGCTTTCTCGATTGCC). Amplicons were confirmed to be the correct size, purified and expanded in a Topo cloning vector (Invitrogen). Six KO and three KO2 clones were purified and sent for Sanger sequencing (Eurofins Genomics).

### Transfection of siRNA

siRNAs to HVEM and PTP1B (Ambion, ThermoFisher Scientific) were reverse transfected with Lipofectamine 2000 to a final concentration of 20 nM and left for 48 h. The Silencer Select negative control siRNA number 1 was used as a negative control (Ambion, ThermoFisher Scientific).

### Quantitation of mRNA by RT-qPCR

Total RNA was isolated using an RNeasy mini kit (Qiagen) then DNase I (Invitrogen) treated according to the manufacturers protocol. cDNA synthesis was performed with Superscript III (Invitrogen) and random hexamers according to the manufacturer’s instructions. Quantitative polymerase chain reaction (qPCR) was performed with the MESA BLUE qPCR kit for SYBR assay (Eurogentec) on a LightCycler96 system (Roche) using primers shown in Table S1.

### Immunofluorescence and confocal microscopy

Cells for immunofluorescence were grown on coverslips and fixed with 4% paraformaldehyde in PBS for 20 min at room temperature, followed by permeabilisation with 0.5% Triton-X100 for 10 min. Fixed cells were blocked by incubation in PBS with 10% NCS for 20 min, before the addition of primary antibody in PBS with 10% NCS, and a further 30-min incubation. After extensive washing with PBS, the appropriate Alexafluor conjugated secondary antibody was added in PBS with 10% NCS and incubated for a further 30 min. The coverslips were washed extensively in PBS and mounted in Mowiol containing DAPI. Images were acquired using a Nikon A1 confocal microscope and processed using ImageJ software [46].

## Acknowledgements

The authors thank Carlos Maluquer de Motes, Richard Roller, David Johnson and Colin Crump for reagents used in this study. We are also grateful to Gill Wallis and the BIAFC staff at University of Surrey for their assistance with flow cytometry and cell sorting.

---

**Figure S1.**
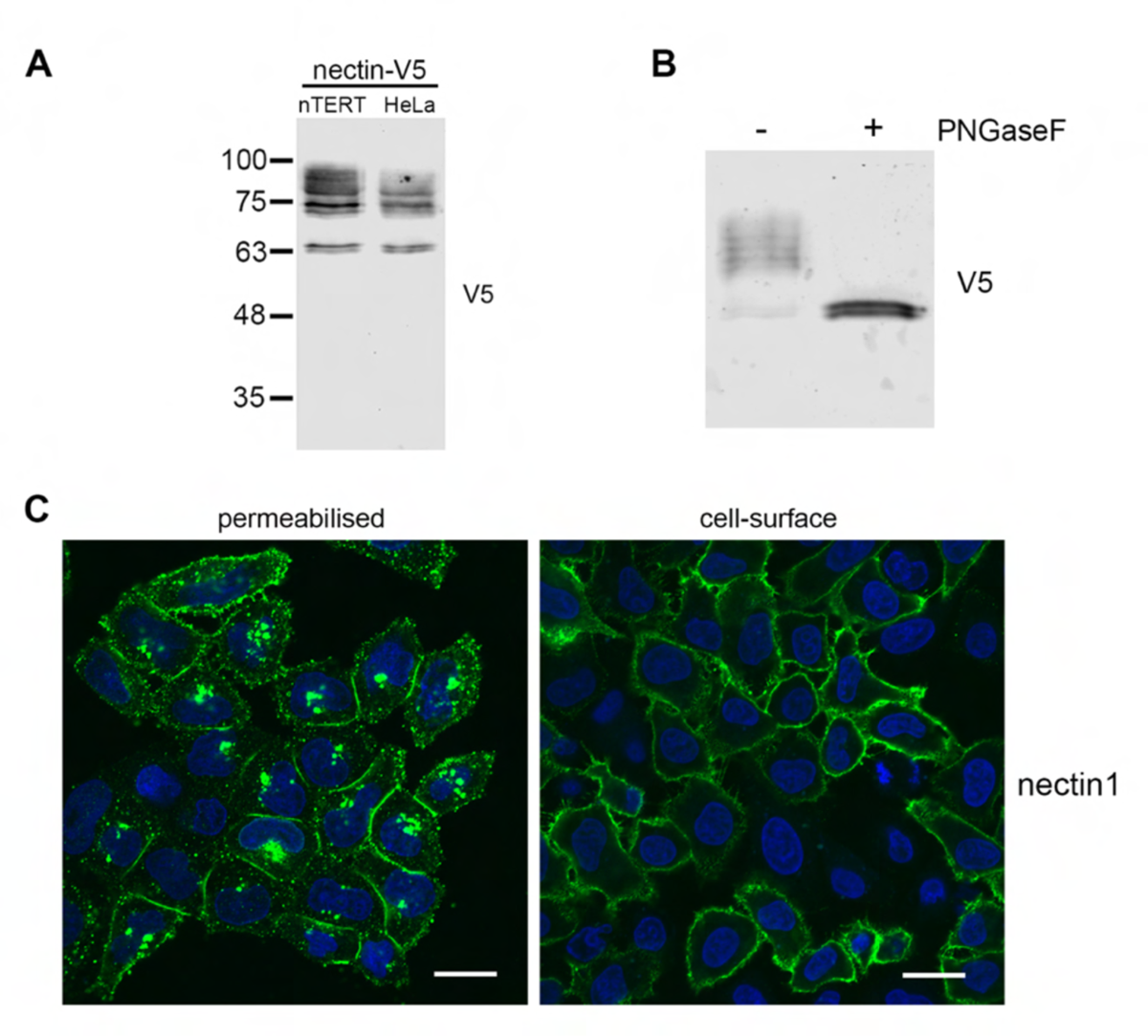
**(A)** nTERT and HeLa cells were transfected with plasmid expressing nectin1V5, harvested at 16 h and analysed by SDS-PAGE and Western blotting for V5. **(B)** As for (**A**), but nectin1-V5 transfected nTERT cells were harvested and lysates subjected to deglycosylation with PNGaseF prior to analysing by SDS-PAGE and Western blotting. (**C**) nTERT cells grown on coverslips were transfected with nectin1-V5-expressing plasmid. Sixteen hours later, cells were either cell-surface stained with antibody to the extracellular domain of nectin1 prior to fixation, or fixed and permeabilised followed by staining with the same antibody (green). Nuclei were stained with DAPI (blue). Scale bar = 20 μm.

**Figure S2.**
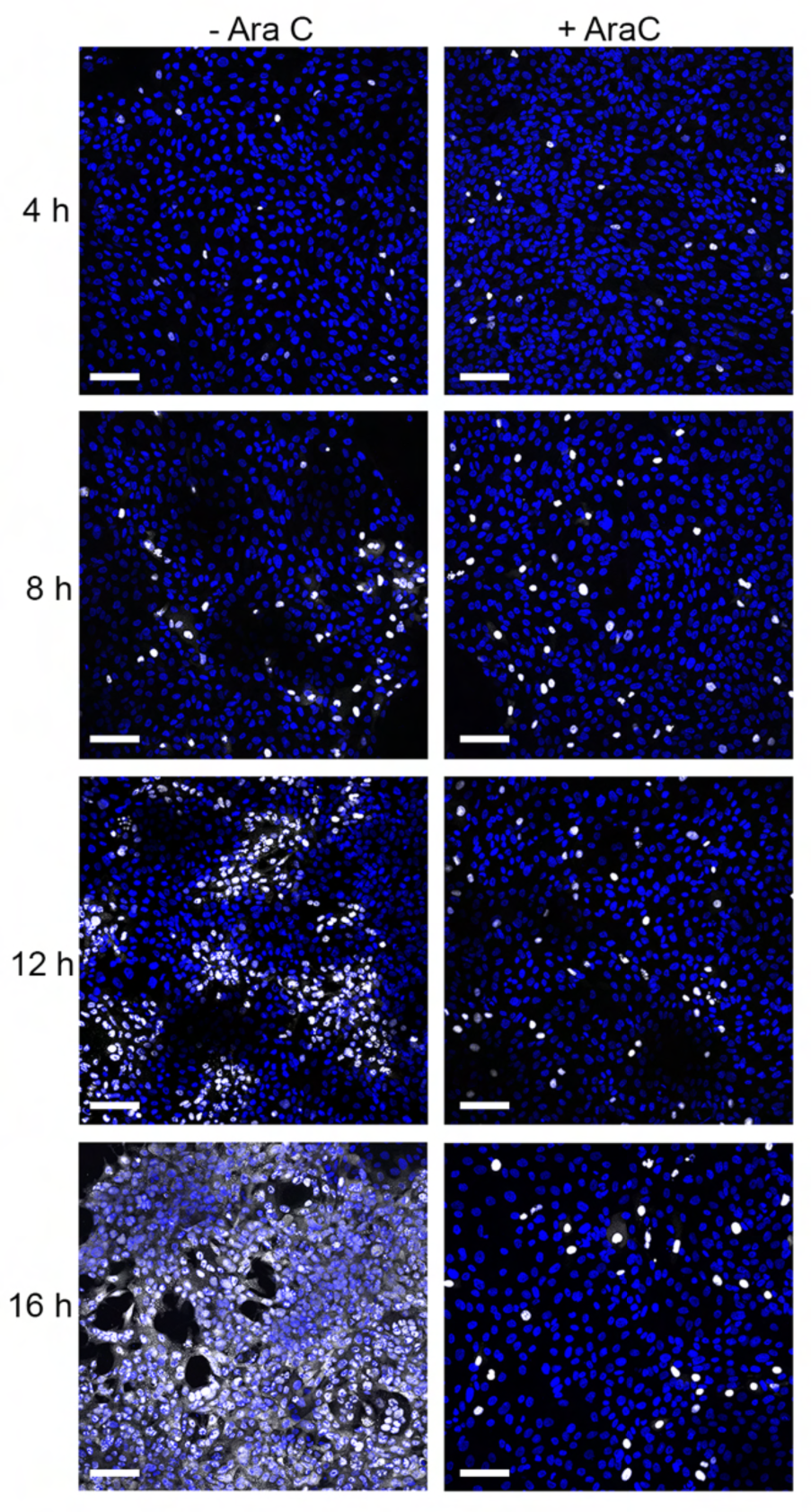
Confluent nTERT cells were infected with Sc16 at MOI 0.01 in the absence or presence of 100 ng/ml AraC, fixed and permeabilised at the indicated times and stained for ICP4 (white) and nuclei were stained with DAPI (blue). Scale bar = 100μm.

**Figure S3.**
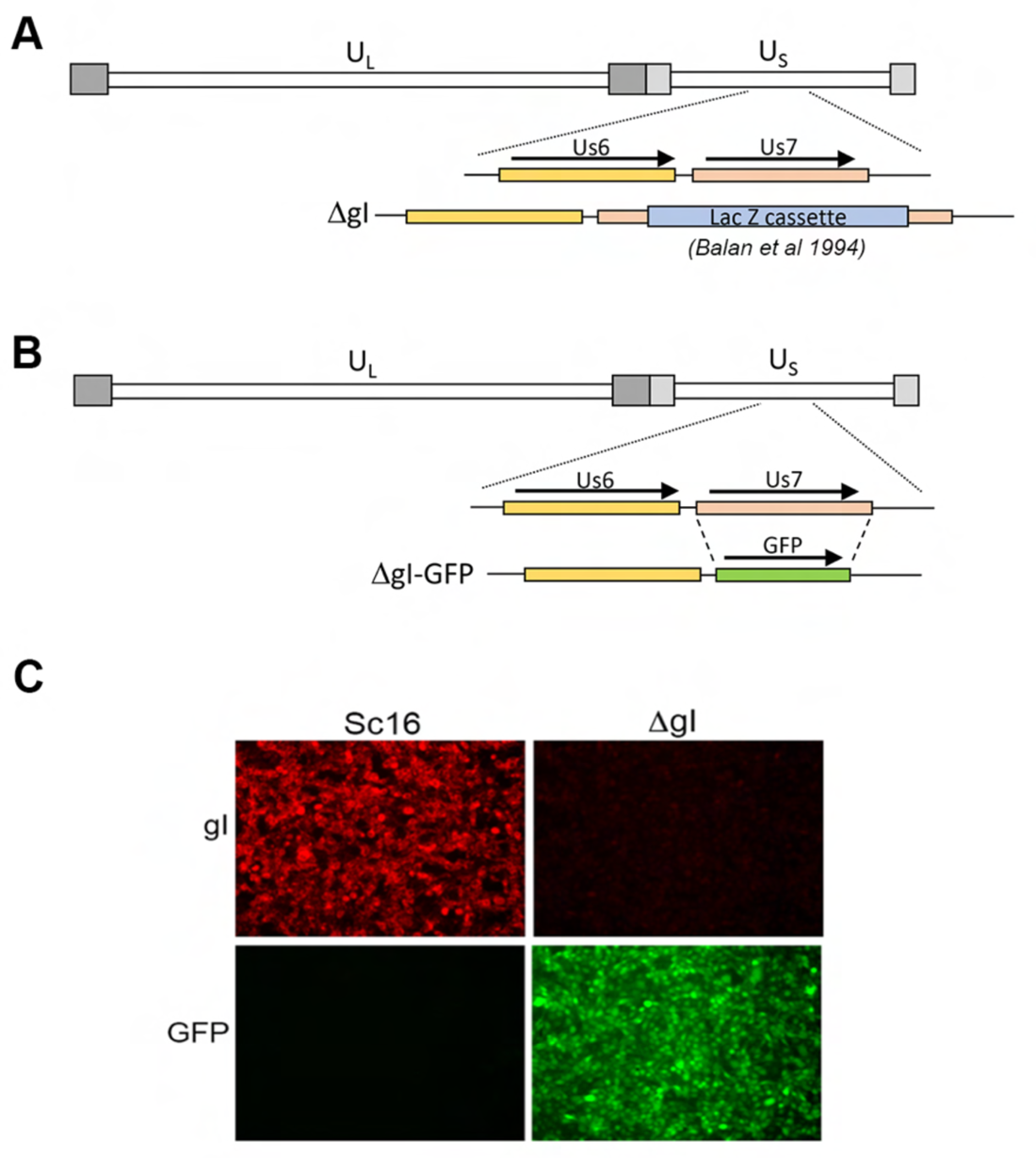
**(A)** Line drawing of the Sc16 *Δ*gI genome that was first described in [15]. **(B)** Line drawing of Sc16 *Δ*gI-GFP constructed here by replacing the gI gene Us7 with the open reading frame for GFP using homologous recombination as shown. The proximity of the gD-encoding gene Us6 is also indicated. **(C)** Vero cells infected with Sc16 or *Δ*gI-GFP were fixed and permeabilised at 16 hours, and imaged for gI (stained in red) and GFP (green).

**Figure S4.**
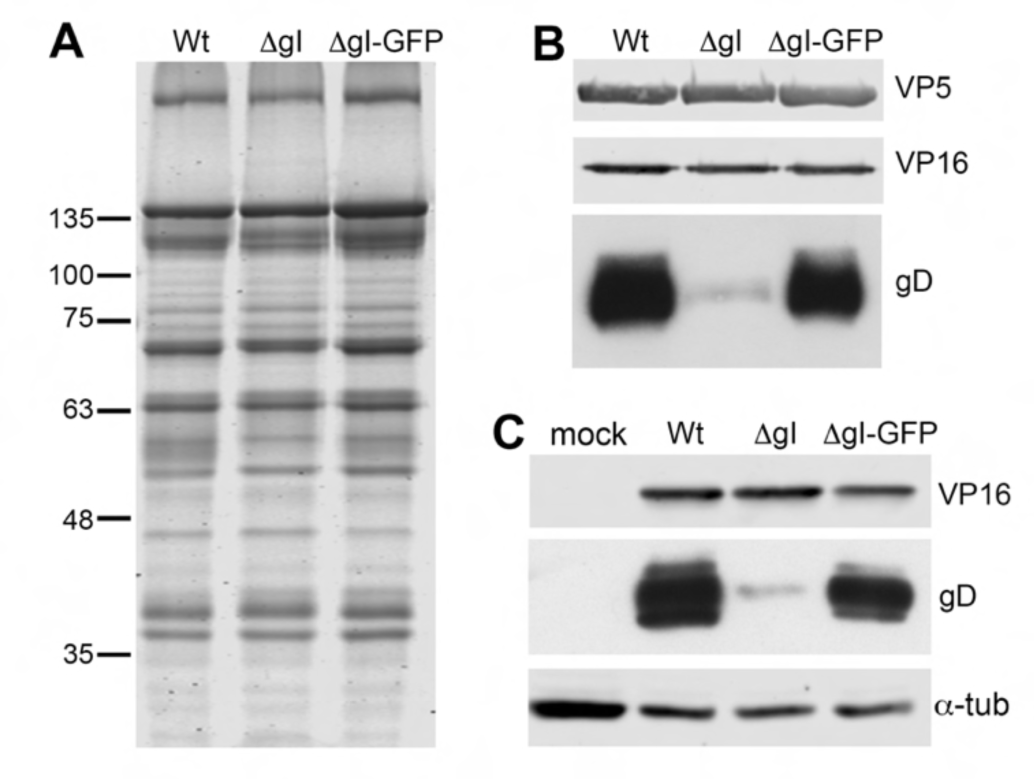
(A) & (B) Extracellular virions were purified from HaCaT cells infected with Sc16 (Wt), *Δ*gI or *Δ*gIGFP, and analysed by SDS-PAGE followed by staining with Coomassie blue **(A)** or Western blotting for the major capsid protein VP5, VP16 and gD **(B)**. **(C)** nTERT cells infected with Wt, *Δ*gI or *Δ*gIGFP viruses were harvested at 16 hours and analysed by SDS-PAGE and Western blotting for VP16, gD and *α*-tubulin.

**Figure S5.**
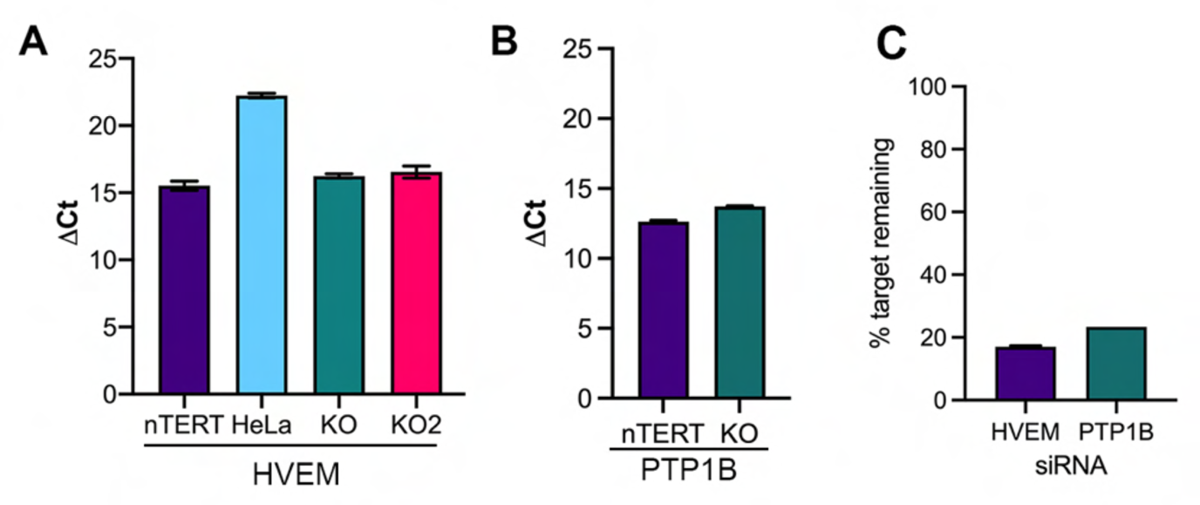
**(A)** Total RNA was isolated from HeLa, nTERT and KO cells, and subjected to RT-qPCR using primers for HVEM. Results are presented as *Δ*Ct measurement using 18s as reference (mean±SEM, *n* = 3). **(B)** Total RNA was isolated from nTERT and KO cells and subjected to RT-qPCR using primers for PTP1B. Results are presented as *Δ*Ct measurement using 18s as reference (mean±SEM, *n* = 3). **(C)** KO cells were transfected with control, HVEM or PTP1B siRNA, total RNA harvested after 48 h and RT-qPCR performed. Level of mRNA in siRNA transfected cells is presented as the percentage of mRNA in control siRNA transfected cells.

**Table S1.**
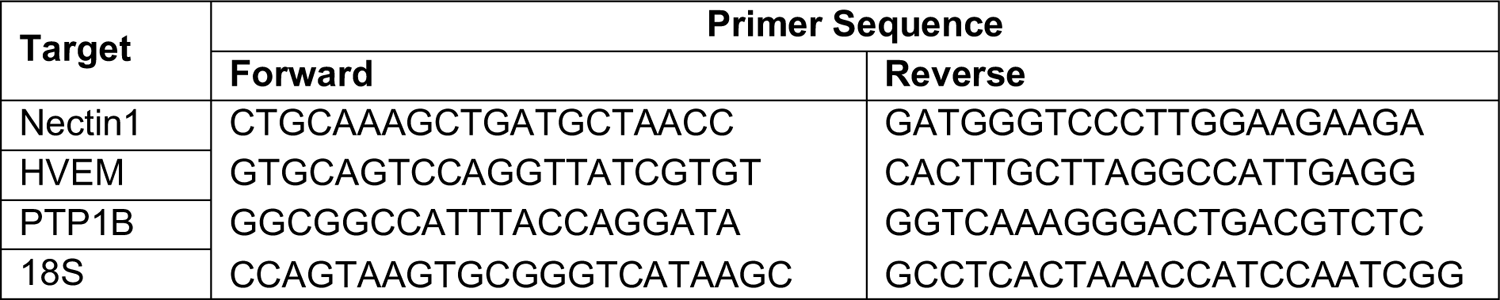
qPCR primers used for measuring siRNA knockdown efficiency.

## References

1. Kramer T, Enquist LW. Directional spread of alphaherpesviruses in the nervous system. Viruses. 2013;5(2):678–707. Epub 2013/02/26. doi: 10.3390/v5020678. PubMed PMID: 23435239; PubMed Central PMCID: PMCPMC3640521.

2. Boelsma E, Verhoeven MC, Ponec M. Reconstruction of a human skin equivalent using a spontaneously transformed keratinocyte cell line (HaCaT). J Invest Dermatol. 1999;112(4):489–98. Epub 1999/04/14. doi: 10.1046/j.1523-1747.1999.00545.x. PubMed PMID: 10201534.

3. Dickson MA, Hahn WC, Ino Y, Ronfard V, Wu JY, Weinberg RA, et al. Human keratinocytes that express hTERT and also bypass a p16(INK4a)-enforced mechanism that limits life span become immortal yet retain normal growth and differentiation characteristics. Mol Cell Biol. 2000;20(4):1436–47. PubMed PMID: 10648628.

4. Smits JPH, Niehues H, Rikken G, van Vlijmen-Willems I, van de Zande G, Zeeuwen P, et al. Immortalized N/TERT keratinocytes as an alternative cell source in 3D human epidermal models. Scientific reports. 2017;7(1):11838. Epub 2017/09/21. doi: 10.1038/s41598-017-12041-y. PubMed PMID: 28928444; PubMed Central PMCID: PMCPMC5605545.

5. Sayers CL, Elliott G. Herpes Simplex Virus 1 Enters Human Keratinocytes by a Nectin-1-Dependent, Rapid Plasma Membrane Fusion Pathway That Functions at Low Temperature. J Virol. 2016;90(22):10379–89. Epub 2016/10/30. doi: 10.1128/JVI.01582-16. PubMed PMID: 27630229; PubMed Central PMCID: PMCPMC5105671.

6. Eisenberg RJ, Atanasiu D, Cairns TM, Gallagher JR, Krummenacher C, Cohen GH. Herpes virus fusion and entry: a story with many characters. Viruses. 2012;4(5):800–32. Epub 2012/07/04. doi: 10.3390/v4050800. PubMed PMID: 22754650; PubMed Central PMCID: PMCPMC3386629.

7. Geraghty RJ, Krummenacher C, Cohen GH, Eisenberg RJ, Spear PG. Entry of alphaherpesviruses mediated by poliovirus receptor-related protein 1 and poliovirus receptor. Science. 1998;280(5369):1618-20. PubMed PMID: 9616127.

8. Montgomery RI, Warner MS, Lum BJ, Spear PG. Herpes simplex virus-1 entry into cells mediated by a novel member of the TNF/NGF receptor family. Cell. 1996;87(3):427–36. PubMed PMID: 8898196.

9. Marsters SA, Ayres TM, Skubatch M, Gray CL, Rothe M, Ashkenazi A. Herpesvirus entry mediator, a member of the tumor necrosis factor receptor (TNFR) family, interacts with members of the TNFR-associated factor family and activates the transcription factors NF-kappaB and AP-1. J Biol Chem. 1997;272(22):14029–32. Epub 1997/05/30. PubMed PMID: 9162022.

10. Huber MT, Wisner TW, Hegde NR, Goldsmith KA, Rauch DA, Roller RJ, et al. Herpes simplex virus with highly reduced gD levels can efficiently enter and spread between human keratinocytes. J Virol. 2001;75(21):10309–18. Epub 2001/10/03. doi: 10.1128/JVI.75.21.10309-10318.2001. PubMed PMID: 11581399; PubMed Central PMCID: PMC114605.

11. Petermann P, Thier K, Rahn E, Rixon FJ, Bloch W, Ozcelik S, et al. Entry mechanisms of herpes simplex virus 1 into murine epidermis: involvement of nectin-1 and herpesvirus entry mediator as cellular receptors. J Virol. 2015;89(1):262–74. Epub 2014/10/17. doi: 10.1128/JVI.02917-14. PubMed PMID: 25320325.

12. Jones J, Depledge DP, Breuer J, Ebert-Keel K, Elliott G. Genetic and phenotypic intrastrain variation in herpes simplex virus type 1 Glasgow strain 17 syn+-derived viruses. J Gen Virol. 2019;100(12):1701–13. Epub 2019/10/30. doi: 10.1099/jgv.0.001343. PubMed PMID: 31661047.

13. Sattentau QJ. The direct passage of animal viruses between cells. Current opinion in virology. 2011;1(5):396–402. Epub 2012/03/24. doi: 10.1016/j.coviro.2011.09.004. PubMed PMID: 22440841.

14. Dingwell KS, Brunetti CR, Hendricks RL, Tang Q, Tang M, Rainbow AJ, et al. Herpes simplex virus glycoproteins E and I facilitate cell-to-cell spread in vivo and across junctions of cultured cells. J Virol. 1994;68(2):834–45. PubMed PMID: 8289387.

15. Balan P, Davis-Poynter N, Bell S, Atkinson H, Browne H, Minson T. An analysis of the in vitro and in vivo phenotypes of mutants of herpes simplex virus type 1 lacking glycoproteins gG, gE, gI or the putative gJ. J Gen Virol. 1994;75 1245–58. PubMed PMID: 8207391.

16. Cocchi F, Menotti L, Dubreuil P, Lopez M, Campadelli-Fiume G. Cell-to-cell spread of wild-type herpes simplex virus type 1, but not of syncytial strains, is mediated by the immunoglobulin-like receptors that mediate virion entry, nectin1 (PRR1/HveC/HIgR) and nectin2 (PRR2/HveB). J Virol. 2000;74(8):3909-17. Epub 2000/03/23. doi: 10.1128/jvi.74.8.3909-3917.2000. PubMed PMID: 10729168; PubMed Central PMCID: PMCPMC111902.

17. Carmichael JC, Yokota H, Craven RC, Schmitt A, Wills JW. The HSV-1 mechanisms of cell-to-cell spread and fusion are critically dependent on host PTP1B. PLoS Pathog. 2018;14(5):e1007054. Epub 2018/05/10. doi: 10.1371/journal.ppat.1007054. PubMed PMID: 29742155; PubMed Central PMCID: PMCPMC5962101.

18. Lachmann RH, Sadarangani M, Atkinson HR, Efstathiou S. An analysis of herpes simplex virus gene expression during latency establishment and reactivation. J Gen Virol. 1999;80 (Pt 5):1271–82. Epub 1999/06/04. doi: 10.1099/0022-1317-80-5-1271. PubMed PMID: 10355774.

19. Sayers CL, Elliott G. Herpes Simplex Virus 1 Enters Human Keratinocytes by a Nectin-1-Dependent, Rapid Plasma Membrane Fusion Pathway That Functions at Low Temperature. Journal of Virology. 2016;90(22):10379–89. doi: 10.1128/jvi.01582-16.

20. Roller RJ, Zhou Y, Schnetzer R, Ferguson J, DeSalvo D. Herpes simplex virus type 1 U(L)34 gene product is required for viral envelopment. J Virol. 2000;74(1):117–29. Epub 1999/12/10. doi: 10.1128/jvi.74.1.117-129.2000. PubMed PMID: 10590098; PubMed Central PMCID: PMCPMC111520.

21. Jolly C, Kashefi K, Hollinshead M, Sattentau QJ. HIV-1 cell to cell transfer across an Env-induced, actin-dependent synapse. The Journal of experimental medicine. 2004;199(2):283–93. Epub 2004/01/22. doi: 10.1084/jem.20030648. PubMed PMID: 14734528; PubMed Central PMCID: PMC2211771.

22. Cudmore S, Reckmann I, Griffiths G, Way M. Vaccinia virus: a model system for actin-membrane interactions. J Cell Sci. 1996;109(Pt 7):1739–47.

23. Hong JS, Ju HJ. The Plant Cellular Systems for Plant Virus Movement. Plant Pathol J. 2017;33(3):213–28. Epub 2017/06/09. doi: 10.5423/PPJ.RW.09.2016.0198. PubMed PMID: 28592941; PubMed Central PMCID: PMCPMC5461041.

24. Cifuentes-Munoz N, Dutch RE, Cattaneo R. Direct cell-to-cell transmission of respiratory viruses: The fast lanes. PLoS Pathog. 2018;14(6):e1007015. Epub 2018/06/29. doi: 10.1371/journal.ppat.1007015. PubMed PMID: 29953542; PubMed Central PMCID: PMCPMC6023113.

25. Singh BK, Pfaller CK, Cattaneo R, Sinn PL. Measles Virus Ribonucleoprotein Complexes Rapidly Spread across Well-Differentiated Primary Human Airway Epithelial Cells along F-Actin Rings. mBio. 2019;10(6). Epub 2019/11/28. doi: 10.1128/mBio.02434-19. PubMed PMID: 31772054; PubMed Central PMCID: PMCPMC6879720.

26. Lawrence DM, Patterson CE, Gales TL, D’Orazio JL, Vaughn MM, Rall GF. Measles virus spread between neurons requires cell contact but not CD46 expression, syncytium formation, or extracellular virus production. J Virol. 2000;74(4):1908–18. Epub 2000/01/22. doi: 10.1128/jvi.74.4.1908-1918.2000. PubMed PMID: 10644364; PubMed Central PMCID: PMCPMC111669.

27. Panasiuk M, Rychlowski M, Derewonko N, Bienkowska-Szewczyk K. Tunneling Nanotubes as a Novel Route of Cell-to-Cell Spread of Herpesviruses. J Virol. 2018;92(10). Epub 2018/03/02. doi: 10.1128/JVI.00090-18. PubMed PMID: 29491165; PubMed Central PMCID: PMCPMC5923070.

28. Dingwell KS, Johnson DC. The herpes simplex virus gE-gI complex facilitates cell-to-cell spread and binds to components of cell junctions. J Virol. 1998;72(11):8933–42. PubMed PMID: 9765438.

29. Dingwell KS, Doering LC, Johnson DC. Glycoproteins E and I facilitate neuron-to-neuron spread of herpes simplex virus. J Virol. 1995;69(11):7087–98. Epub 1995/11/01. doi: 10.1128/JVI.69.11.7087-7098.1995. PubMed PMID: 7474128; PubMed Central PMCID: PMCPMC189628.

30. Howard PW, Wright CC, Howard T, Johnson DC. Herpes simplex virus gE/gI extracellular domains promote axonal transport and spread from neurons to epithelial cells. J Virol. 2014;88(19):11178–86. Epub 2014/07/18. doi: 10.1128/JVI.01627-14. PubMed PMID: 25031334; PubMed Central PMCID: PMCPMC4178820.

31. Johnson DC, Webb M, Wisner TW, Brunetti C. Herpes simplex virus gE/gI sorts nascent virions to epithelial cell junctions, promoting virus spread. J Virol. 2001;75(2):821–33. PubMed PMID: 11134295.

32. McGraw HM, Awasthi S, Wojcechowskyj JA, Friedman HM. Anterograde spread of herpes simplex virus type 1 requires glycoprotein E and glycoprotein I but not Us9. J Virol. 2009;83(17):8315–26. Epub 2009/07/03. doi: 10.1128/JVI.00633-09. PubMed PMID: 19570876; PubMed Central PMCID: PMCPMC2738194.

33. DuRaine G, Wisner TW, Howard P, Williams M, Johnson DC. Herpes Simplex Virus gE/gI and US9 Promote both Envelopment and Sorting of Virus Particles in the Cytoplasm of Neurons, Two Processes That Precede Anterograde Transport in Axons. J Virol. 2017;91(11). Epub 2017/03/24. doi: 10.1128/JVI.00050-17. PubMed PMID: 28331094; PubMed Central PMCID: PMCPMC5432863.

34. Scherer J, Hogue IB, Yaffe ZA, Tanneti NS, Winer BY, Vershinin M, et al. A kinesin-3 recruitment complex facilitates axonal sorting of enveloped alpha herpesvirus capsids. PLoS Pathog. 2020;16(1):e1007985. Epub 2020/01/30. doi: 10.1371/journal.ppat.1007985. PubMed PMID: 31995633; PubMed Central PMCID: PMCPMC7010296.

35. Diwaker D, Murray JW, Barnes J, Wolkoff AW, Wilson DW. Deletion of the Pseudorabies Virus gE/gI-US9p complex disrupts kinesin KIF1A and KIF5C recruitment during egress, and alters the properties of microtubule-dependent transport in vitro. PLoS Pathog. 2020;16(6):e1008597. Epub 2020/06/09. doi: 10.1371/journal.ppat.1008597. PubMed PMID: 32511265; PubMed Central PMCID: PMCPMC7302734.

36. Wilkinson GW, Davison AJ, Tomasec P, Fielding CA, Aicheler R, Murrell I, et al. Human cytomegalovirus: taking the strain. Medical microbiology and immunology. 2015;204(3):273–84. Epub 2015/04/22. doi: 10.1007/s00430-015-0411-4. PubMed PMID: 25894764; PubMed Central PMCID: PMCPMC4439430.

37. Oliver SL, Yang E, Arvin AM. Varicella-Zoster Virus Glycoproteins: Entry, Replication, and Pathogenesis. Curr Clin Microbiol Rep. 2016;3(4):204–15. Epub 2017/04/04. doi: 10.1007/s40588-016-0044-4. PubMed PMID: 28367398; PubMed Central PMCID: PMCPMC5373811.

38. Warner MS, Geraghty RJ, Martinez WM, Montgomery RI, Whitbeck JC, Xu R, et al. A cell surface protein with herpesvirus entry activity (HveB) confers susceptibility to infection by mutants of herpes simplex virus type 1, herpes simplex virus type 2, and pseudorabies virus. Virology. 1998;246(1):179–89. doi: 10.1006/viro.1998.9218. PubMed PMID: 9657005.

39. Satoh T, Arii J, Suenaga T, Wang J, Kogure A, Uehori J, et al. PILRalpha is a herpes simplex virus-1 entry coreceptor that associates with glycoprotein B. Cell. 2008;132(6):935–44. doi: 10.1016/j.cell.2008.01.043. PubMed PMID: 18358807; PubMed Central PMCID: PMC2394663.

40. Hill TJ, Field HJ, Blyth WA. Acute and recurrent infection with herpes simplex virus in the mouse: a model for studying latency and recurrent disease. J Gen Virol. 1975;28(3):341–53. Epub 1975/09/01. doi: 10.1099/0022-1317-28-3-341. PubMed PMID: 170376.

41. Elliott G, O’Hare P. Live-cell analysis of a green fluorescent protein-tagged herpes simplex virus infection. J Virol. 1999;73:4110–9.

42. Hutchinson I, Whiteley A, Browne H, Elliott G. Sequential Localization of Two Herpes Simplex Virus Tegument Proteins to Punctate Nuclear Dots Adjacent to ICP0 Domains. J Virol. 2002;76(20):10365–73.

43. Carter GC, Rodger G, Murphy BJ, Law M, Krauss O, Hollinshead M, et al. Vaccinia virus cores are transported on microtubules. J Gen Virol. 2003;84(Pt 9):2443–58. PubMed PMID: 12917466.

44. Russell T, Bleasdale B, Hollinshead M, Elliott G. Qualitative Differences in Capsidless L-Particles Released as a By-Product of Bovine Herpesvirus 1 and Herpes Simplex Virus 1 Infections. J Virol. 2018;92(22). Epub 2018/09/07. doi: 10.1128/JVI.01259-18. PubMed PMID: 30185590; PubMed Central PMCID: PMCPMC6206470.

45. Russell T, Samolej J, Hollinshead M, Smith GL, Kite J, Elliott G. Novel role for ESCRT-III component CHMP4C in the integrity of the endocytic network utilized for herpes simplex virus envelopment. bioRxiv. 2020: 2020.08.19.258558. doi: 10.1101/2020.08.19.258558.

46. Schneider CA, Rasband WS, Eliceiri KW. NIH Image to ImageJ: 25 years of image analysis. Nature methods. 2012;9(7):671–5. Epub 2012/08/30. PubMed PMID: 22930834; PubMed Central PMCID: PMCPmc5554542.

